# Microtubules form by progressively faster tubulin accretion, not by nucleation-elongation

**DOI:** 10.1101/545236

**Authors:** Luke M. Rice, Michelle Moritz, David A. Agard

## Abstract

Microtubules are dynamic polymers that play fundamental roles in all eukaryotes. Despite their importance, how new microtubules form is poorly understood. Textbooks have focused on variations of a nucleation-elongation mechanism in which monomers rapidly equilibrate with an unstable oligomer (nucleus) that limits the rate of polymer formation; once formed, the polymer then elongates efficiently from this nucleus by monomer addition. Such models faithfully describe actin assembly, but they fail to account for how more complex polymers like hollow microtubules assemble. Here we articulate a new model for microtubule formation that has three key features: i) microtubules initiate via rectangular, sheet-like structures which grow faster the larger they become; ii) the dominant pathway proceeds via accretion, stepwise addition of longitudinal or lateral layers; iii) a ‘straightening penalty’ to account for the energetic cost of tubulin’s curved-to-straight conformational transition. This model can quantitatively fit experimental assembly data, providing new insights into biochemical determinants and assembly pathways for microtubule nucleation.

## Introduction

The microtubule cytoskeleton plays essential roles organizing the interior of eukaryotic cells: microtubules can act as “tracks” for motor-based transport, they form the basis of the mitotic spindle that mediates proper segregation of chromosomes during cell division, they help define cell polarization, and more (Barlan and Gelfand, 2017; Desai and Mitchison, 1997; Prosser and Pelletier, 2017). To fulfill these many roles, microtubules must assemble into a multitude of highly dynamic and spatially diverse networks. The dynamic instability of microtubules (Mitchison and Kirschner, 1984), which describes the switching of individual polymers between phases of growing and shrinking, allows wholesale reorganization of the microtubule network to occur rapidly and also provides a mechanism to efficiently search space (Holy and Leibler, 1994). Indeed, numerous regulatory factors allow cells to control how fast microtubules grow and how frequently they switch from growing to shrinking (catastrophe) or from shrinking back to growing (rescue) (Akhmanova and Steinmetz, 2015), and also to control the creation of new microtubules (Kollman et al., 2011; Roostalu and Surrey, 2017; Wieczorek et al., 2015), a process called nucleation.

Microtubules are hollow, cylindrical polymers formed from αβ-tubulin heterodimers that interact with each other in two ways: stronger head-to-tail (longitudinal) interactions between αβ-tubulins make up the straight protofilaments, and weaker side-to-side (lateral) interactions hold protofilaments together. The mechanisms underlying the dynamic instability of existing microtubules are increasingly well understood (Alushin et al., 2014; Driver et al., 2017; Duellberg et al., 2016; Gardner et al., 2011; Geyer et al., 2015; Grishchuk et al., 2005; Manka and Moores, 2018; Margolin et al., 2012; Piedra et al., 2016; VanBuren et al., 2005; VanBuren et al., 2002; Zhang et al., 2015), but models for how *existing* microtubules grow and shrink cannot specify how *new* microtubules form de novo. Thus, our understanding of the mechanisms that govern spontaneous microtubule formation remains relatively primitive (reviewed in

(Roostalu and Surrey, 2017)). Multiple factors contribute to this difficulty: first, nucleation occurs very rarely and cannot truly be observed microscopically, so measuring it directly is much harder than measuring the growing and shrinking of existing microtubules. Second, the open, tube-like structure of the microtubule poses unique complications. In most organisms, microtubules contain 13 protofilaments, although there are clear examples of specialized microtubules containing 11 or 15 protofilaments (Burton et al., 1975; Chaaban et al., 2018; Chalfie and Thomson, 1982; Davis and Gull, 1983; Kwiatkowska et al., 2006; Saito and Hama, 1982; Tucker et al., 1992). Regardless of protofilament number, the hollow nature of the microtubule means that it takes many more αβ-tubulin subunits to close a tube than it does to make a minimal helical repeat for a simpler, helical polymer like actin. Perhaps not surprisingly given this complexity, there is no consensus about the sequence of intermediates that must be formed (pathway) during the transition from individual subunits to a closed tube. To further complicate matters, the structure of the free heterodimer also changes during assembly, from a bent confirmation in solution to a straight conformation within the final microtubule ((Ayaz et al., 2012; Nawrotek et al., 2011; Rice et al., 2008); reviewed in (Brouhard and Rice, 2018)). Where in the assembly process the straightening takes place, whether it is continuous or step-wise, how much energy is required, and how this obligatory conformational change affects nucleation all remain unknown. In addition to these factors, the GTPase activity required for microtubule catastrophe can also antagonize nucleation by destabilizing intermediate species.

An early study of the kinetics of spontaneous microtubule nucleation, which used solution conditions that suppress catastrophe to simplify the behavior, posited that nucleation proceeded sequentially through two discrete intermediates (Voter and Erickson, 1984). The data were re-analyzed in later studies (Flyvbjerg et al., 1996; Kuchnir Fygenson et al., 1995) that proposed a more complex pathway containing four distinct nucleation intermediates (6-mer, 9-mer, 12-mer, and 15-mer of αβ-tubulin). However, in this later analysis it was not possible to ascribe a biochemical interpretation to the species or rate constants (Flyvbjerg et al., 1996). Thus, the physical-chemical details of the underlying assembly process continue to remain obscure.

In this work, we sought to obtain more explicit insight into the molecular mechanisms that govern spontaneous microtubule assembly. Our primary goal was to explore the simplest biochemical models that could faithfully explain the experimental data. We began from the fundamental modes of αβ-tubulin interactions within a microtubule: a single subunit (heterodimer) can make a longitudinal interaction, a lateral interaction or make both simultaneously (corner interaction). Starting with explicit simulations of the biochemical reactions that can occur for all possible αβ-tubulin oligomers up to 12-mers, we observed that a small subset of all possible oligomers became dominant. These dominant oligomers were “rectangular”, meaning that they maximized the number of longitudinal and lateral contacts such that there were no unfilled corners. The dominance of a small subset of species allowed us to develop a simpler, approximate model based almost exclusively on rectangular oligomers. This simplification allowed the model to access arbitrarily large oligomers in a way that retained a connection to the fundamental biochemical interaction modes of the individual subunits.

This approximate but biochemically faithful model provides a new way to view microtubule nucleation. Rather than proceeding via a distinct rare, rate limiting “nucleus”, the model indicates that microtubule assembly involves the continuous accretion of αβ-tubulins into a growing 2D lattice. Like crystallization processes, the larger the assembly, the faster it grows. Our model reveals assembly pathways for forming a new microtubule and provides new insights into how exogeneous nucleators and other regulatory factors can accelerate microtubule formation.

## Results

### Rectangular species dominate the assembly pathway

To obtain biochemical insight into possible pathways for spontaneous microtubule assembly, we constructed an explicit model that includes all non-redundant tubulin oligomers from dimers through dodecamers (Fig. 1A shows the intermediates through hexamers). Here non-redundant means that we only consider a single exemplar N-mer for each unique combination of longitudinal and lateral contacts; statistical weighting of rate constants ensures that this exemplar oligomer faithfully represents the ensemble of species it represents (see example in Sup. Fig. 1). We defined the biochemical pathways that connect different oligomers (arrows in Fig. 1A) by assuming that oligomers grow and shrink via addition and loss of individual αβ-tubulin subunits. It was not practical to explore larger oligomers in this formulation because of an explosion in the number of species and reactions (reaching dodecamers requires ~200 distinct species and ~600 reactions) (Fig. 1F).

**Figure 1.**
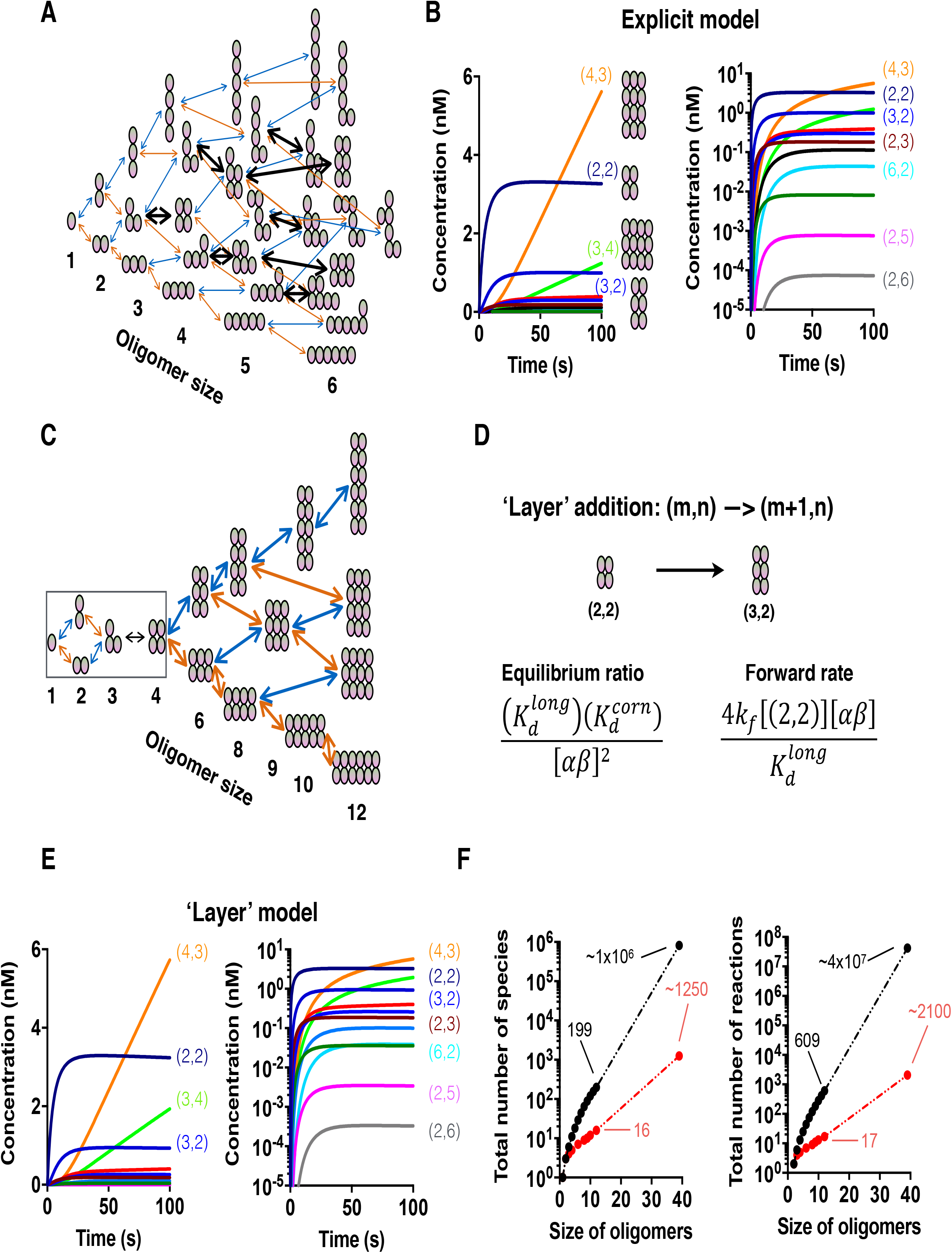
An explicit biochemical model for early steps in spontaneous microtubule nucleation. **A** Cartoon of the model. Individual αβ-tubulin heterodimers (ovals) can associate longitudinally (blue arrows) or laterally (orange arrows). As oligomers grow in size, they can present sites where an associating tubulin would simultaneously make longitudinal and lateral interactions (“corner”; bold black arrows). The resulting collection of species and reactions was implemented in BerkeleyMadonna (Macey et al., 2009) for simulations. Species up to 6-mers are shown, but the model includes species up to 12-mers. Rapid growth of the number of oligomers and reactions with size made it impractical to consider even larger oligomers. **B** (left) Plot showing that ‘rectangular’ species dominate the simulation. The four dominant species are shown, with values in parentheses indicating the height and width for the oligomers. (right) The same data plotted on a logarithmic concentration scale. Other rectangular oligomers are much less common. Non-rectangular oligomers are not shown, but are also much rarer than the dominant oligomers shown. **C** A simplified ‘layer growth’ model that only considers rectangular oligomers. Arrows indicate longitudinal (blue) or lateral (orange) layer growth. The resulting collection of species and reactions was implemented in BerkeleyMadonna for simulations (see Methods) **D** Cartoon of a ‘layer’ addition reaction with associated rate constants. The equilibrium ratio and rate for layer addition can be defined in terms of underlying longitudinal, lateral, and corner affinities and a single on rate constant. An effective equilibrium constant and an approximation of the effective forward rate constant for this layer addition are shown. See Methods for derivations. **E** Similar to **B** but showing results from simulations of the simplified layer model. This much simpler model with many fewer species and reactions quantitatively reproduces results from the explicit model. **F** (left) Plot of the species count for all possible oligomers (black) and for the subset of rectangular oligomers (red). The extrapolation to a 40-mer emphasizes the magnitude of the simplification. (right) Similar plot but for the number of biochemical reactions.

We implemented the differential equations describing this discrete assembly scheme in Berkeley Madonna (Marcoline et al., 2020) to simulate the assembly pathway and kinetics. The only free parameters in this model are the on rate constant kf and the affinities for the three different αβ–tubulin interaction modes (longitudinal, lateral, and corner (simultaneous longitudinal+lateral interaction (Fig 1D). As is common, kf was assumed (here we chose 10^6^ M^-1^s^-1^ based on experimental observations and consistent with work from others (Coombes et al., 2013; Margolin et al., 2012; McIntosh et al., 2018; VanBuren et al., 2002; Zakharov et al., 2015)). We explored different values for the three parameters, seeking to identify conditions under which some appreciable concentration of dodecamers would form. We were especially interested in the corner affinity, because this is the interaction that would provide the driving force for microtubule elongation. A relatively tight corner affinity (K_d_^corner^ < 0.2 μM; Sup. Fig. 2) was required for appreciable flux into the largest oligomers. With high-affinity corner interactions, the simulations predominantly populated a small number of ‘rectangular’ oligomers, defined as oligomers wherein (i) every subunit makes at least one longitudinal and one lateral contact to another, and (ii) there are no ‘open’ corner sites (Fig. 1B). More explicitly, what happens in the simulations is that formation of an initial 2×2 oligomer forms relatively quickly. The next larger species containing an additional ‘singleton’ tubulin are poorly populated because of the weak longitudinal and lateral affinities, but those species also present an open, high-affinity corner site. Filling this open site stabilizes the first subunit and in the case of a 2×2 start, fills the next longitudinal or lateral layer resulting in a 3×2 or 2×3 assembly, respectively. On larger assemblies, additional corners are vacant after the first, and these fill rapidly, essentially “zipping up” the layer. The net consequence is that once initiated, layer completion occurs efficiently, explaining why rectangular species dominate.

Only considering rectangular oligomers greatly reduces the number of species and reactions required to reach much larger oligomers in simulations (e.g. Fig. 1C,F). However, in such an approximation the rates of adding or losing a layer become context dependent, and depend on the height and width of the oligomer. Using the steady-state approximation, we were able to derive a general recursive expression for the effective ‘forward’ rate of adding of a longitudinal or lateral layer (see Methods). An example of adding a longitudinal layer to a 2×2 oligomer is illustrated in Fig. 1D. Considering the layer addition as a sequence of individual steps also allowed us to derive an effective forward rate as well as a ‘layer equilibrium ratio’, which describes the expected relative populations of oligomers that differ by a single layer (Methods). Importantly, in these expressions both the forward rate and equilibrium ratio for layer addition are defined in terms of the biochemical properties of individual subunits (longitudinal, lateral and corner affinities/rates). Consequently, and in contrast to what occurs with phenomenological models, the reduction in the number of species and reactions does not come at the expense of losing a connection to the fundamental biochemical behavior. For the 12-mer simulated explicitly above, the layer model provides a >10-fold reduction in species and >35-fold reduction in number of reactions (Fig. 1F) (these reduction factors increase dramatically for larger oligomer sizes).

We implemented the reduced, layer-based model into Berkeley Madonna to test if it could recapitulate the results from our explicit simulations when using the same input parameters for subunit affinities. Despite the dramatic reduction in complexity, and with the exception of very low-abundance intermediate species, the simplified layer-based simulations quantitatively reproduced the oligomer concentrations and the kinetics of oligomer formation observed with the explicit model (compare Fig. 1E to 1B). These initial benchmarking layer simulations only considered oligomers up to dodecamers so that we could make a direct comparison with the explicit model. In the next section, we expand the model to include the much larger oligomers required to depict microtubule assembly.

### A simplified accretion pathway for spontaneous microtubule assembly

We wrote a custom program to expand the layer model to arbitrary sized rectangular oligomers. Whenever an oligomer gained a 13^th^ protofilament, we converted it into a microtubule. No additional penalty for seam formation was included. We assumed that once formed, microtubules elongate by capturing αβ-tubulin subunits one at a time in the open, pseudo-helical ‘corner’ sites on either end. In order to minimize the number of adjustable parameters, the same rate constants were used for events at either the plus- or minus-ends. These additional simplifying assumptions allow the model to simulate spontaneous microtubule assembly and the ensuing depletion of free αβ-tubulin subunitsin a relatively natural way, without having to explicitly specify a particular nucleus or assume a preferred assembly pathway.

We began testing our model by fitting it to experimental assembly curves that we measured by light scattering (Fig. 2A). The model was able to fit individual curves quite well (Fig. 2B), with minor discrepancies evident at low concentration. The optimized affinities differed somewhat from fit to fit, but shared common features: tight corner affinities (K_d_^corn^ < 100 nM), weaker longitudinal affinities (K_d_^long^ ~5-15 mM), and lateral affinities around 10-fold weaker than longitudinal affinities (Fig. 2B). The longitudinal affinity we obtained is in the range of that obtained from biochemical models for microtubule growing and shrinking (e.g.(Gardner et al., 2011; Kim and Rice, 2019; Piedra et al., 2016; VanBuren et al., 2002)). The corner and lateral affinities are both about 10-fold stronger than in those models, a difference that might reflect the different buffer conditions (high glycerol concentration) in ‘nucleation’ assays relative to ‘dynamics’ assays. The corner affinity is much stronger than the product of longitudinal and lateral affinities, which is expected because the individual longitudinal and lateral affinities represent a balance between (among other things) favorable protein:protein contacts and the unfavorable entropic cost of losing rotational and translational degrees of freedom -- when making a corner interaction the longitudinal and lateral interfaces both contribute favorably while the entropic cost remains approximately the same (see also (Erickson and Pantaloni, 1981)). Fits to an independent set of assembly curves (Sup. Fig. 3A) yielded similar affinities (Sup. Fig. 3B). We found it encouraging that the model could fit individual assembly curves, but to be convincing a single parameter set should be able to fit all the assembly curves simultaneously. Thus, we turned to global fitting where we attempted to fit the set of assembly curves with one common parameter set.

**Figure 2.**
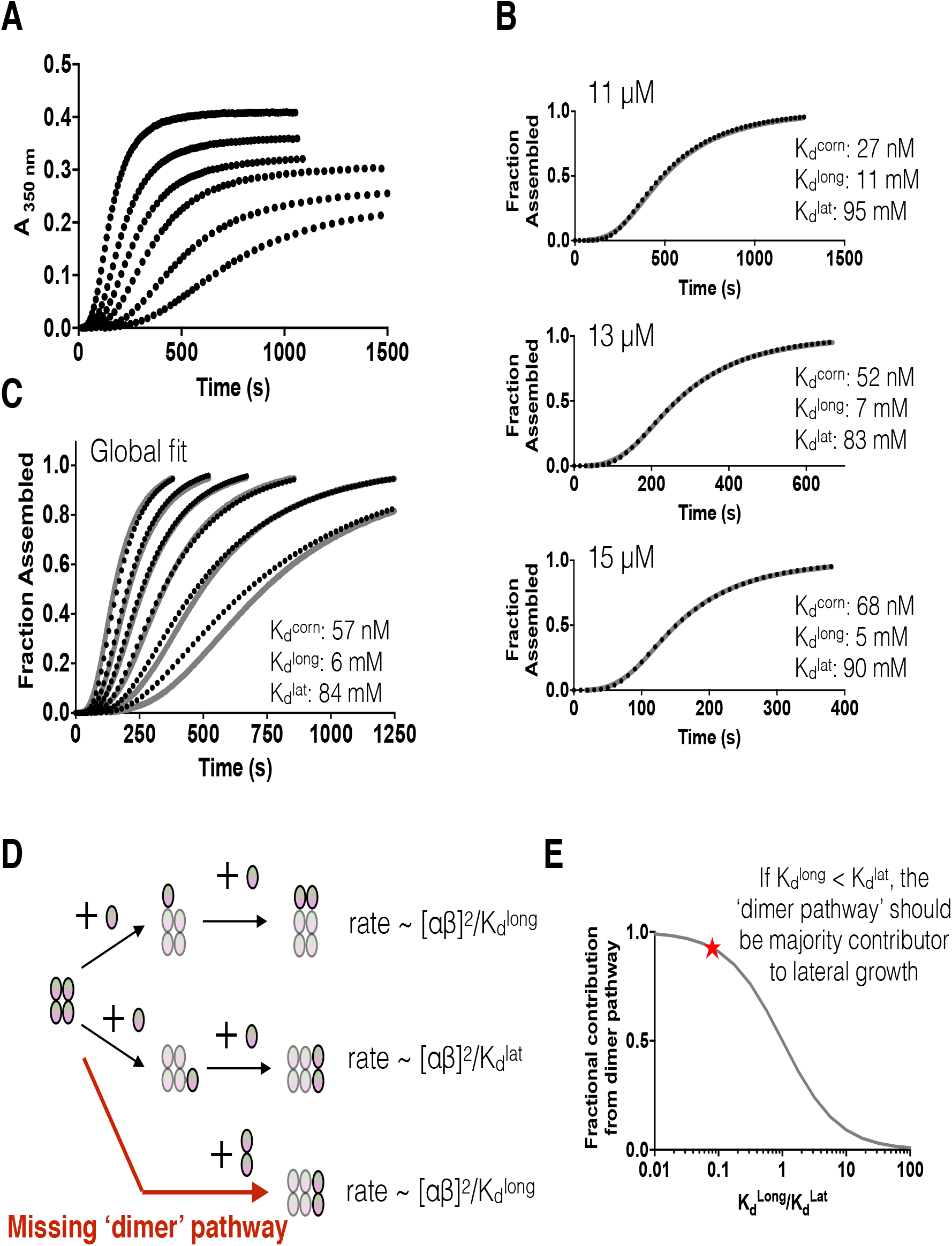
Expansion of the layer model to arbitrarily large oligomers for simulating spontaneous microtubule nucleation. Oligomers are allowed to become arbitrarily tall. We assumed that when oligomers reached a width of 13, they ‘closed’ into a tube and elongated by monomer addition. **A** Spontaneous microtubule nucleation kinetics measured by turbidity (A350 nm) at multiple concentrations of αβ-tubulin (10, 11, 12, 13, 14, 15 μM). **B** Fits of the model (black dotted line) to individual assembly curves (solid grey line) at the indicated concentration. Fits to individual curves are good. Inset text shows the resulting longitudinal, lateral, and ‘corner’ (simultaneous longitudinal and lateral interactions) affinities, which differ somewhat from run to run. **C** Global fit of the model (black dashed lines) to the full set of assembly data (grey lines). As evidenced by the poor fit at low concentration, the model underestimates the concentration-dependence of the progress curves. **D** Cartoon illustrating that longitudinal dimers of αβ-tubulin should contribute to lateral growth. The associated forward rates for the indicated reactions are shown. **E** The relative contribution of ‘monomer’ and ‘dimer’ initiation pathways can be estimated as indicated based on the affinities of longitudinal and lateral interactions. Plot predicting the fractional contribution of a ‘dimer’ pathway for lateral growth in terms of the relative strength of longitudinal and lateral interactions. The red star indicates the ratio obtained from the fit shown in **B**. Those parameters predict that the ‘dimer’ pathway should dominate the assembly, but it is not accounted for in our model.

In a global fit to the set of assembly curves (Fig. 2C), a limitation of the model became apparent: even though each individual concentration could be well fit (Fig. 2B), the model could not capture the correct behavior over the broad concentration range measured (Fig. 2C). The model appeared to fit the alternative dataset somewhat better (Sup. Fig. 3C), but it was still apparent that the model was underestimating the concentration-dependence of the assembly curves; an outlier curve also limited the concentration range available for fitting. Because all layer transition rates scale with the chosen on-rate constant (see derivations in Methods), changing the assumed on-rate does not improve the global fits. The observed discrepancy between model and experiment suggests that the model fails to capture some important aspect of assembly. In re-examining our assumptions, we realized that it was too restrictive to consider only monomer addition and loss as the ‘elemental’ reactions within the layer assembly pathway. Indeed, a simple calculation demonstrates that any model having these relative lateral and longitudinal affinities must also include a ‘dimer pathway’ (Fig. 2D). Compared to the ‘two step’ monomer-based pathway in which a first monomer associates weakly and only becomes ‘locked in’ by a subsequent corner association (Fig. 2D, middle path), when the longitudinal affinity is stronger than the lateral one as it is for microtubules it is more efficient to initiate a lateral layer by adding a pre-formed longitudinal dimer (Fig. 2D, lower path). This is true even though longitudinal dimers are present at very low concentrations (roughly 1 part in a 1000 or ~10 nM) for the parameters we used. Our ‘monomer only’ model global fit required a longitudinal affinity ~10-fold stronger than the lateral one, but under those conditions additions of longitudinal dimers should actually account for ~90% of the lateral layer initiations (Fig. 2E). Thus, despite appearing reasonable, the fits do not make biochemical sense because the model fails to account for a dimer pathway that has the potential to dominate the flux through oligomers. It has been suggested that oligomer associations may be important during spontaneous microtubule assembly (Erickson and Pantaloni, 1981; Mozziconacci et al., 2008; Voter and Erickson, 1984), but to our knowledge these interactions have not previously been implemented into kinetic models.

### Incorporating a dimer pathway fails to recapitulate experimental data without invoking a switch in lateral affinity

The cartoons and calculations presented in Fig. 2D,E illustrate how the rate of initiating a new lateral layer will depend on the concentration of longitudinal dimers of αβ-tubulin. By contrast, once the slow step of *initiating* a new layer has occurred, *zipping up* via addition into favorable ‘corner’ (longitudinal + lateral) sites should occur almost exclusively via αβ-tubulin ‘monomers’ because these monomers far exceed longitudinal dimers in number/concentration. By a similar logic, it is also not necessary to include a dimer pathway for initiating a longitudinal layer: lateral dimers are very rare, and the ‘upper’ αβ-tubulin of a longitudinal dimer provides no increased stabilization as a longitudinal layer builds up.

Incorporating a parallel, dimer-based pathway for initiating lateral layers (Sup. Fig. 4) requires that at each step we specify (i) the concentration of longitudinal dimers, (ii) an association rate constant for the longitudinal dimers adding to a complete layer, and (iii) a dissociation rate constant for the lateral ‘unbinding’ of a longitudinal dimer. The concentration of longitudinal dimers is easily calculated given the concentration of monomers and the equilibrium constant for longitudinal interactions. Because off-rate constants tend to vary over a much larger range than on-rate constants, for simplicity we used the same, roughly diffusion limited association rate constant as we do for monomers. We derived a formula that expresses the equilibrium constant for lateral association of a longitudinal dimer solely in terms of biochemical properties of the monomers (see Methods). Thus, we were able to parameterize the ‘dimer pathway’ purely in terms of the fundamental monomer properties.

We first fit the dimer model to individual assembly curves, as we did for the monomer model (Fig. 3A). The longitudinal and corner affinities for these dimer model fits were comparable to those obtained from fits of the monomer model. The lateral affinity (~25-30 M in the dimer model) was ~300-fold weaker than the in the monomer model (~100 mM), reflecting a greatly diminished contribution of the monomer pathway for lateral initiation. Although it was predicted to be required biochemically, including this dimer pathway actually degraded the ability of the model to recapitulate the shape of individual assembly curves compared to the monomer-only fits: the simulated assembly curves displayed a less pronounced lag phase and also appeared less cooperative (shallower sigmoid transition; compare Fig. 3A to Fig. 2B). Global fits of the dimer model to the set of assembly curves were also markedly poorer than for the monomer model (Fig. 3B). Thus, a more complete biochemical model for the assembly process paradoxically degraded its predictive power.

**Figure 3.**
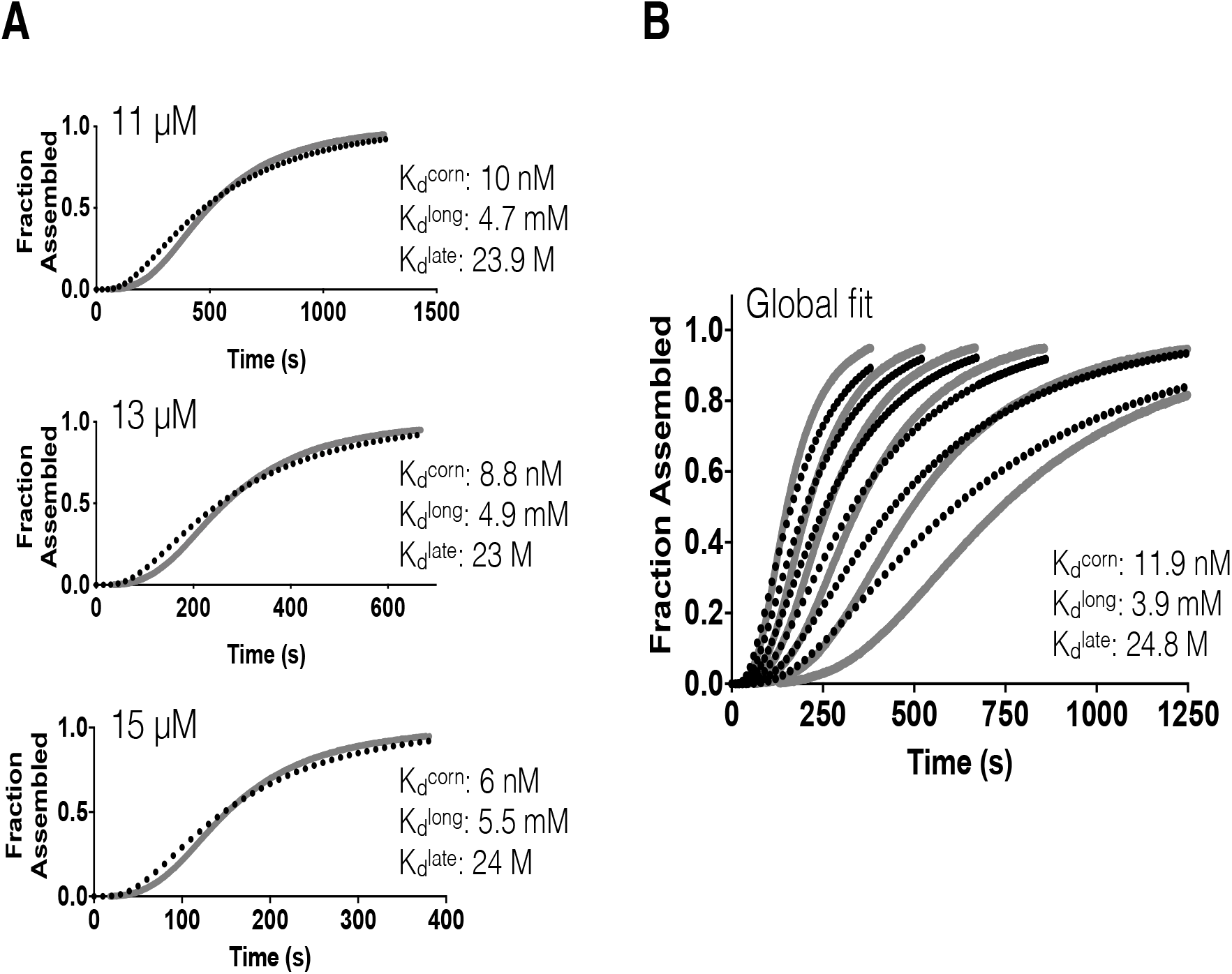
Incorporating a dimer pathway for lateral growth into the model **A** Fits of the new dimer model (black dotted line) to individual assembly curves (solid grey line) at the indicated concentrations. Fits to individual curves are worse than for the monomer only model (compare to Fig. 2B). Inset text shows the resulting longitudinal, lateral, and ‘corner’ (simultaneous longitudinal and lateral interactions) affinities; the corner affinity varies almost two-fold over for 11 and 15 μM fits. The lateral affinity is ~200-fold weaker than for ‘monomer model’ fits, reflecting the dominance of the dimer pathway in this model. **B** Global fit of the model (black dashed lines) to the full set of assembly data (grey lines). The fit is poor, with the model predicting too fast assembly at all concentrations.

Why was the monomer-only model able to fit the observed assembly curves so much better than a model incorporating a dimer pathway? The presence of a dimer pathway makes the effective rates of lateral and longitudinal growth much more comparable to each other (Fig. 2D, compare top and bottom rate expressions), whereas in the monomer-only model there is significantly more contrast between the rates for longitudinal and lateral growth. We speculated that the otherwise more accurate monomer+dimer model must still be missing a critical feature that acts to increase the difference between longitudinal and lateral growth.

What might be missing? Not having accounted for the effects of conformational changes in αβ-tubulin that occur during microtubule formation (Ayaz et al., 2012; Nawrotek et al., 2011; Pecqueur et al., 2012; Wang and Nogales, 2005) is a strong candidate. Indeed, it is now well established that unpolymerized αβ-tubulin subunits adopt a ‘curved’ conformation in solution and that the microtubule lattice itself acts as an allosteric activator to ‘straighten’ αβ-tubulin during the assembly process (Ayaz et al., 2012; Buey et al., 2006; Nawrotek et al., 2011; Rice et al., 2008). That is, αβ-tubulin becomes progressively straighter as it becomes increasingly surrounded by lateral neighbors (reviewed in (Brouhard and Rice, 2018)) (Fig. 4A). This straightening-induced conformational strain acts to weaken lateral interactions in assemblies that require more straightening. To explore these ideas in the context of our model, we introduced a very simple lateral penalty to mimic the greater strain intrinsic to straighter conformations. We assumed that two-wide assemblies could remain curved, whereas wider assemblies would be forced to straighten, making the lateral interactions after the first pairing in a given ‘row’ X-fold weaker.

**Figure 4.**
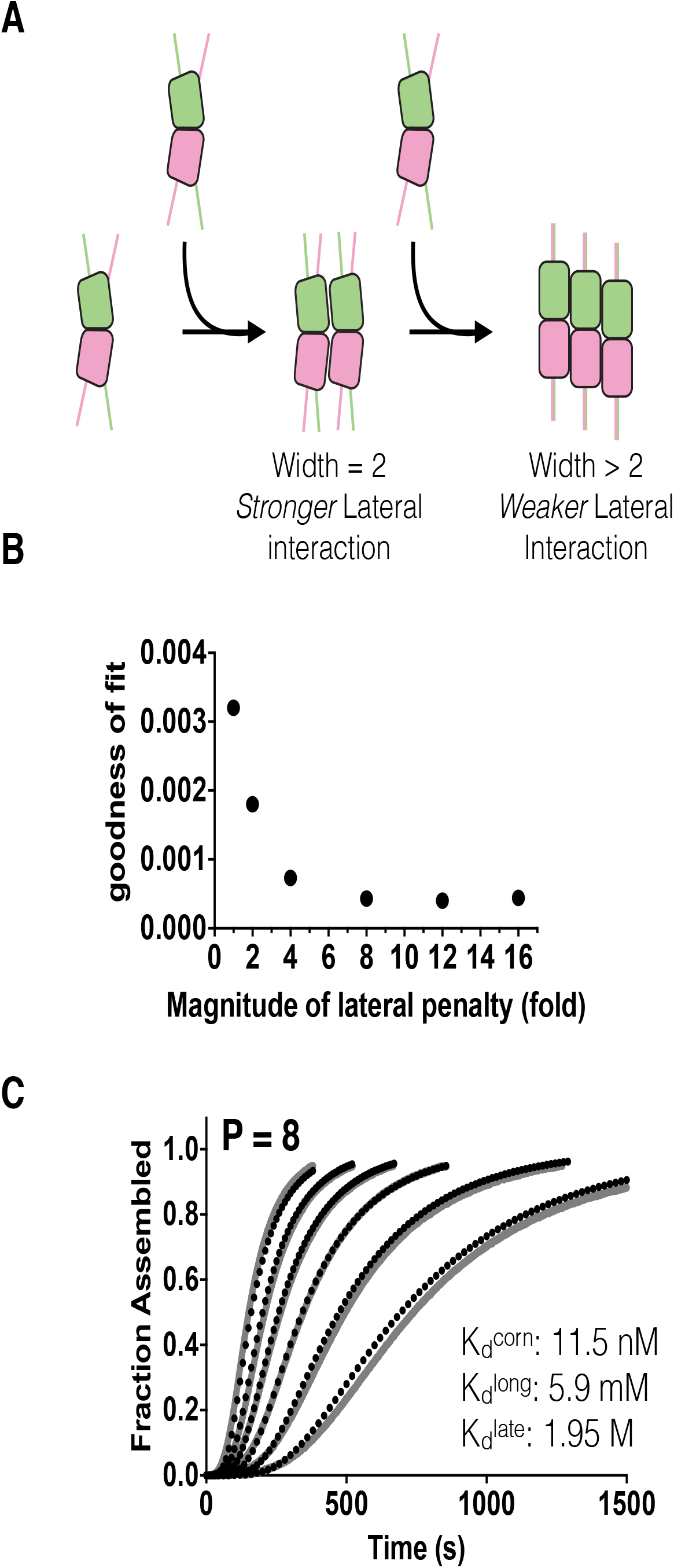
A ‘penalty’ that mimics the cost of αβ-tubulin straightening markedly improves the performance of the dimer model **A** Cartoons illustrating αβ-tubulin straightening, which is thought to occur progressively with width (the number of lateral neighbors). To model this in a very simple way, we introduced a penalty to make lateral interactions weaker for the 3^rd^ and successive lateral interactions. **B** Performance of the ‘dimer with penalty’ model as a function of penalty strength. The improved goodness of fit from the global optimization is not very sensitive to the magnitude of the penalty, as long as the penalty is ~8 or greater. **C** Illustration of the model fit to the experimental data for a penalty value of 8. Inset shows fitted affinities (the corner affinity given here is for width=2 interactions, i.e. before application of the penalty).

We fit the monomer+dimer+penalty model to the data as before, testing a range of penalty magnitudes (from 1 to 16 in Fig. 4B). The global fit to the experimental curves improved as the magnitude of the lateral penalty increased, with the best fits reached for penalty values greater than 4. Thus, penalizing lateral growth was sufficient to ‘correct’ the deficiencies of the simple dimer pathway, resulting in even better fits (Fig. 4B,C; penalty value = 8,) than for the monomer-only model (see Fig. 3B,C). Beyond a value of 4, the improvement in global fit only weakly depends on the precise value of the penalty (Fig. 4B). Fits to the alternative dataset yielded similar but not identical affinities (Sup. Fig. 5). Other penalty factors gave fits of comparable quality as long as the penalty factor was greater than 4 (Fig. 4B), but yielded different longitudinal and lateral affinities, as expected. Varying individual parameters shows that the fits are sensitive to small (1.5-fold) changes in the parameter values (Sup. Fig. 6). Differences in the fitted parameters for the primary and alternative dataset suggests that within a narrow range there may be some compensation between corner and longitudinal affinities (compare Fig. 4C to Sup. Fig. 5), Collectively, our computational experiments led us to what we believe to be the first simple, biochemically-based model that can accurately recapitulate the observed concentration-dependent kinetics of spontaneous microtubule assembly without having to assume a specific nucleus.

### A kinetically frustrated pathway for spontaneous microtubule assembly

The model developed here provides a biochemical pathway that explains in a new way why the kinetics of microtubule initiation are slow relative to polymer elongation. As a first way to illustrate the assembly pathway that emerges from fitting our model to the experimental data, we plotted the total concentration and average size of intermediate oligomers as a function of time (Fig. 5A). The total concentration of oligomers peaks at around 200 s in the example shown, and the subsequent decrease is a consequence of the formation of microtubules and of the depletion of free αβ-tubulin. The average oligomer size increases gradually over several hundred seconds, reflecting the slow process of accretion and the low likelihood that large layers will be lost in ‘backward’ reactions.

**Figure 5.**
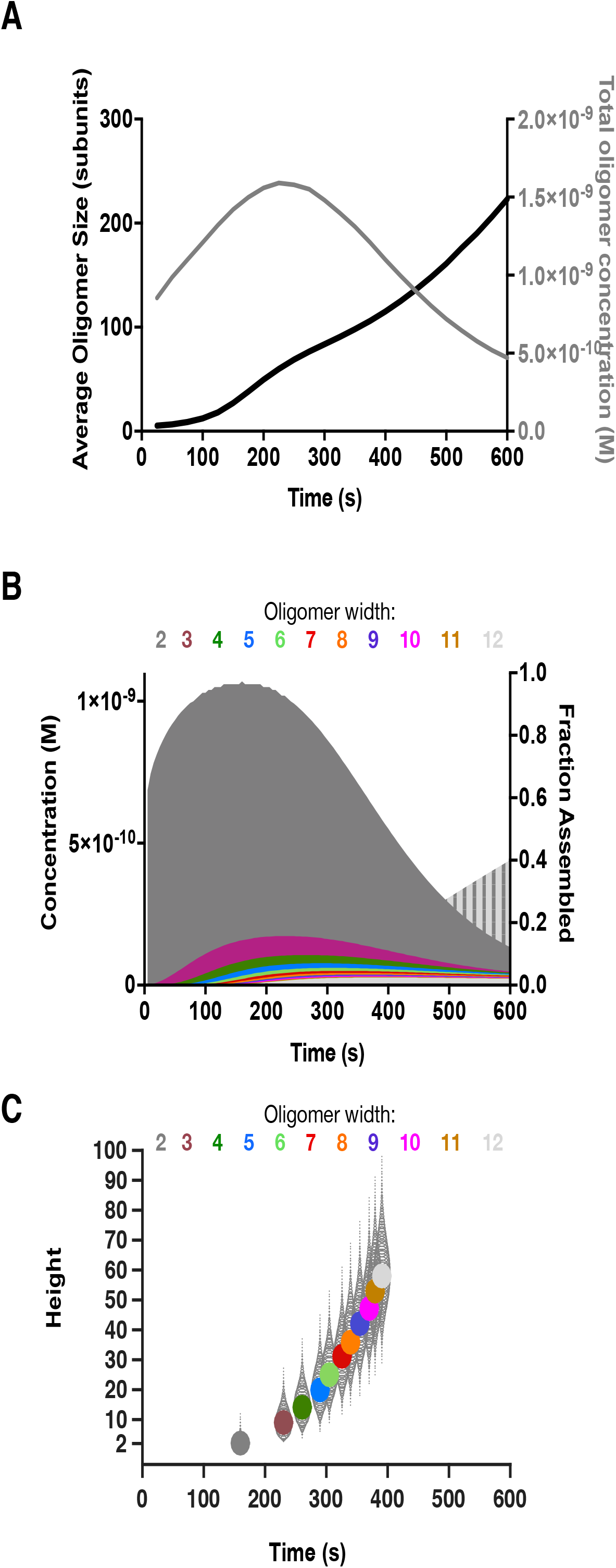
Different ways of representing the predicted pathway for spontaneous microtubule assembly **A** The average size of intermediate oligomers (black curve) increases slowly over time. The total concentration of intermediate oligomers increases and then decreases, an effect that can be explained by depletion of free αβ-tubulin, and the formation of microtubules. **B** Summed concentration for all oligomers of a given width, plotted as a function of time. Oligomers with width=2 dominate. Wider oligomers become progressively rarer and peak at increasingly late times, reflecting the obstacles to lateral growth. The vertically striped light grey curve behind the others shows the extent of total assembly (right y-axis). **C** Distribution of oligomer heights as a function of their width. The most populated height is plotted as a filled circle for each width (different colors), at the time that width peaks. Other height oligomers at a given width are plotted as open circles, with the radius reflecting their concentration relative to the peak species. The time to transition from one width to the next decreases as oligomers get wider and taller, and the spread of oligomer heights increases as oligomers get wider.

The endpoint of microtubule formation – closure of the cylindrical lattice –requires a minimal width oligomer (13 in our model). Thus, we next examined the summed concentration of tubulin oligomers with a given width (lateral extent) as a function of time (Fig. 5B). This analysis shows that the concentration of oligomers decreases as their width increases – there are fewer 3-wide oligomers than 2-wide, fewer 4-wide then 3-wide, etc. (Fig. 5B). This decreasing concentration of wider oligomers happens because the transition from one width to the next accelerates throughout the assembly process. This acceleration can be understood as a statistical/kinetic consequence of the two-dimensional assembly pathway. No matter their height, all oligomers of a given width grow *taller* with the same characteristic rate because they present the same number of ‘landing sites’ for starting a new longitudinal layer. However, taller oligomers grow *wider* faster than shorter oligomers, because their increased height provides more ‘landing sites’ for starting a new lateral layer. Similarly, wider oligomers add longitudinal layers faster than the narrower ones for the same reason.

To provide more granular insight into the nature of the predominant microtubule assembly intermediates, we examined the distribution of oligomer sizes at different widths (Fig. 5C). For each oligomer width, and at the time the concentration of oligomers of that width peaked, we plotted the most populated height (filled circle) and the relative amount of taller or shorter oligomers (grey teardrop/oval shapes, which are actually a set of circles with area scaled proportionally to the concentration). The many parallel pathways in the assembly process leads to an increasing spread in the range of oligomer heights as they get wider. For example, the most populated 50% of all 3-wide oligomers have heights between 7 and 11 (a height range of 5) whereas the most populated 50% of all 12-wide oligomers span heights between 50 and 64 (a height range of 15, three times greater than for 3-wides). The most populated immediate microtubule precursor (12 subunits wide) contains almost 700 tubulins: 58 subunits tall x 12 wide. This is much larger than the number of subunits in the nucleus predicted by phenomenological models (Flyvbjerg et al., 1996; Kuchnir Fygenson et al., 1995). Also note that the rate of layer assembly cannot exceed the rate of elongation. The layer-based assembly model thus provides a new, ensemble-based view of the assembly pathway wherein many oligomers of varying heights contribute appreciably to the formation of new microtubules.

## Discussion

How existing microtubules grow and shrink is increasingly well-understood, but the understanding of how new microtubules form spontaneously – a related but mechanistically distinct process – lags behind (Roostalu and Surrey, 2017). Many discussions of spontaneous microtubule formation have invoked the concept of a ‘critical nucleus’, which is analogous to the transition-state of a chemical reaction (Kollman et al., 2011; Kuchnir Fygenson et al., 1995; Roostalu and Surrey, 2017; Voter and Erickson, 1984). This traditional view of the assembly pathway assumes that (i) the formation and growth of early intermediates is thermodynamically unfavorable, and (ii) above a critical size, oligomer growth by monomer addition becomes favorable and this elongation proceeds at a fixed rate. The polymerization kinetics of relatively simple helical polymers like actin can be well-described by two-step, nucleation-elongation models that invoke a small critical nucleus representing a ‘mini-filament’ (the smallest oligomeric species that presents helical binding sites) (Sept and McCammon, 2001; Tobacman and Korn, 1983). However, the spontaneous assembly kinetics of hollow, cylindrical microtubules do not conform to the simplest two-step, nucleation-elongation kinetics (Flyvbjerg et al., 1996; Kuchnir Fygenson et al., 1995; Voter and Erickson, 1984). Phenomenological multi-step nucleation-elongation mechanisms (Flyvbjerg et al., 1996) can fit assembly curves but fail to provide biochemical insights into the assembly pathway.

Our model provides an alternative and conceptually distinct framework for thinking about the pathway for microtubule formation (Fig. 6), and the biochemical factors that govern the overall rate of assembly. From this comes the understanding that there is not a singular, transition-state-like species (variously estimated as 8-27 tubulins) that dictates the assembly kinetics, but instead what we would traditionally call “nucleation” arises from an ensemble of accretion events involving many species and hundreds of tubulin heterodimers. In what follows, we summarize the main features of the predicted assembly pathway and discuss implications for the multiple ways that a template or other regulatory factor could accelerate the process.

**Figure 6.**
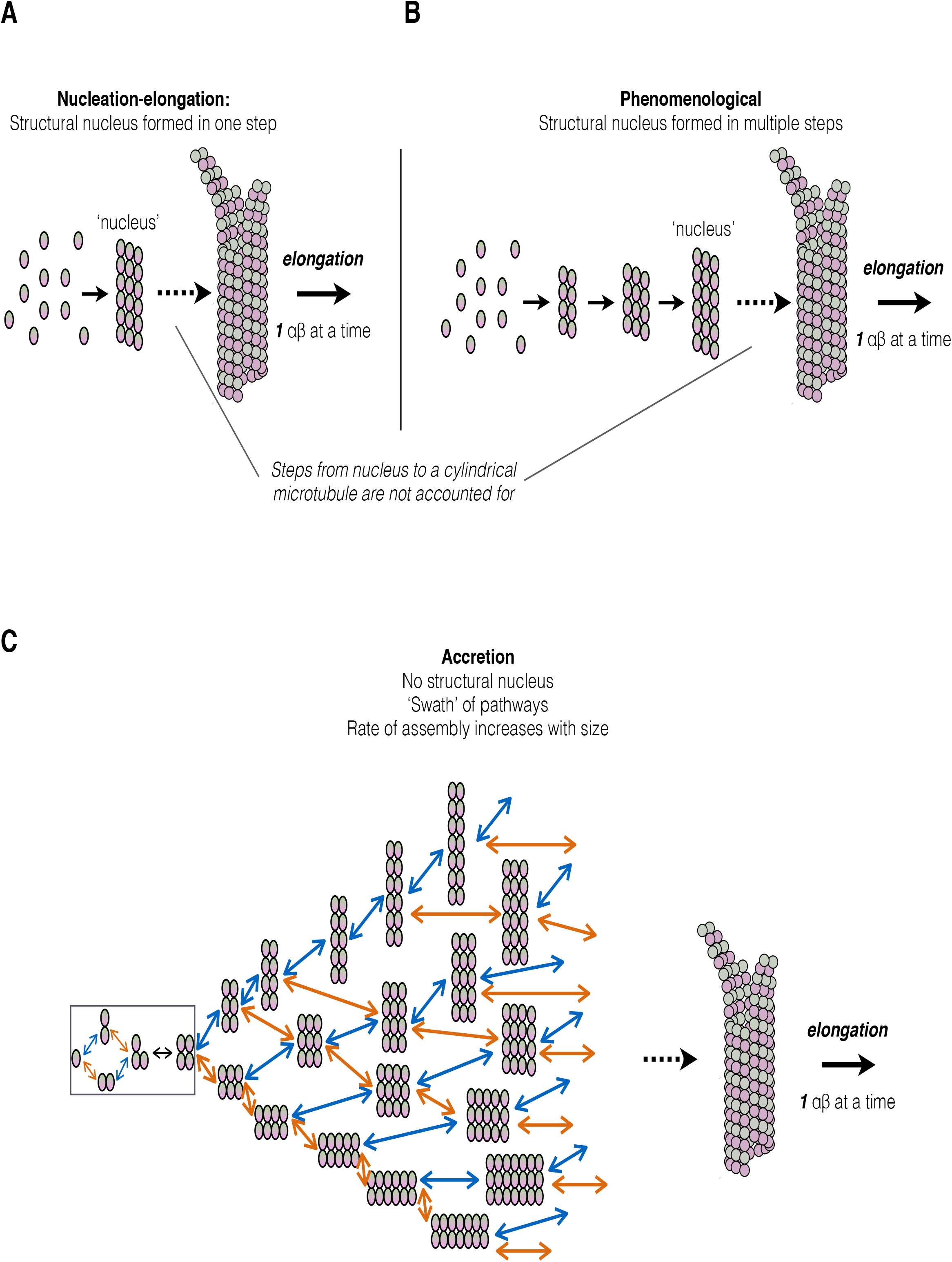
Summary of accretion model and comparison with nucleation-elongation models **A** Cartoon illustrating a classic nucleation-elongation model in which the rate of polymer formation is limited by the concentration of a rare species (the nucleus) that is in a rapid equilibrium with unpolymerized subunits (ovals). Such models cannot account for the additional steps/species in between the nucleus and the formation of the cylindrical microtubule. **B** Cartoon illustrating a phenomenological model (Flyvbjerg et al., I996)_that differs from nucleation-elongation model in that the nucleus is formed in multiple steps. In this model the rate of polymer formation is also limited by the concentration of a rare species (the nucleus), and there is no accounting for the additional steps/species in between the nucleus and the formation of the cylindrical microtubule. **C** Cartoon illustrating the accretion model presented in this paper. In this model there are many pathways to form a cylindrical microtubule, and all ‘layer accretion’ steps are described. In this accretion model, and in contrast with nucleation-elongation and phenomenological models, most of the steps toward polymer formation are energetically favorable but kinetically frustrated by the difficulty of initiating new layers. These barriers to initiation decrease as oligomers become larger, as in crystal growth, and it is the ensemble of pathways and barriers that define the kinetics of microtubule formation.

The layer growth model predicts that a multitude of pathways and intermediates contribute meaningfully to microtubule assembly. Stronger longitudinal interactions cause the dominant rectangular intermediates to be taller than they are wide. Early in the assembly pathway, transitions to larger oligomers become energetically favorable (equilibrium ratio less than one) but still occur slowly because the oligomers are small and because initiating the new layer depends on a weak individual longitudinal or lateral interaction. The rate of transition between oligomers accelerates as oligomers increase in size, not because of any intrinsic changes in the underlying affinities, but because larger oligomers present more sites on which the next layer can initiate. These size-dependent changes in transition rates are a direct consequence of the two-dimensional nature of the assembly pathway. Thus, even though nearly all of the layer additions are energetically ‘downhill’, the repeated difficulty of initiating each new layer contributes a series of progressively smaller delays into the assembly sequence that in combination represent the effective “barrier”. Conformational ‘straightening’ of αβ-tubulins adds an additional kind of frustration that makes lateral growth of early oligomers even harder. Collectively, the ensemble of initiation barriers creates a kind of ‘frustration’ that dictates unique kinetics for spontaneous microtubule assembly.

This layer-based view of the assembly pathway provides new ways for thinking about how regulatory factors might accelerate microtubule initiation. Our modeling indicates that the kinetics of microtubule formation are dictated by the need for *repeated* ‘layer’ transitions that are limited by the weaker lateral affinity and by the straightening penalty (e.g. Fig. 7A). Classical “nucleators” like the γ-tubulin ring complex (γ-TuRC) (Zheng et al., 1995) contain a lateral array 14 γ-tubulins (Fig. 7B) that are thought to form a ‘template’ on which a microtubule can elongate (Consolati et al., 2020; Kollman et al., 2015; Kollman et al., 2011; Liu et al., 2020; Moritz et al., 1995; Wieczorek et al., 2020). We used our model to explore how the potency of templating might depend on the width of the template. We modified our simulations to include various width templating oligomers (Fig. 7C), using a constant total amount of templating ‘monomers’. Note that in these simulations there are progressively lower concentrations of the wider oligomers (e.g. half as many 4-wide oligomers as 2-wide, etc.). The affinity of αβ-tubulins for γ-TuRC is unknown, so we explored a regime where γ:αβ longitudinal interactions are identical to αβ:αβ (Supplemental Fig. 7) and another where γ:αβ longitudinal interactions are weaker than for αβ:αβ (Fig. 7). With the stronger γ:αβ interaction strength, even 2-wide “templates” potently accelerated assembly (Supplemental Fig. 7). This is at odds with experimental data showing that γ-tubulin small complexes (γ-TuSCs), which only contain two γ-tubulins, are much poorer nucleators than γ-TuRCs (Kollman et al., 2015; Oegema et al., 1999). By contrast, when γ:αβ interactions were assumed to be weaker, templating oligomers that were 2 subunits wide only modestly accelerated the predicted assembly kinetics (1.1-fold faster characteristic time for assembly, T0.1) (Fig. 6C,D). A 3-wide oligomer was much more potent (1.7-fold faster) even though in the simulations the concentration of 3-wide oligomers was less than that of the 2-wide oligomer. This strong change in potency occurs because the ‘hardest’ transition in the assembly sequence is from two-wide oligomers to 3-wide oligomers, providing an explanation for why γ-tubulin small complexes (γ-TuSCs) only weakly stimulate assembly (Oegema et al., 1999). A recent study provides experimental support for these prediction by showing that γ:αβ interactions are weaker than αβ:αβ interactions (Thawani et al., 2020) In our simulations, progressively wider templating arrays show even greater potency, emphasizing that the barriers to lateral growth contribute throughout the entire assembly pathway.

**Figure 7.**
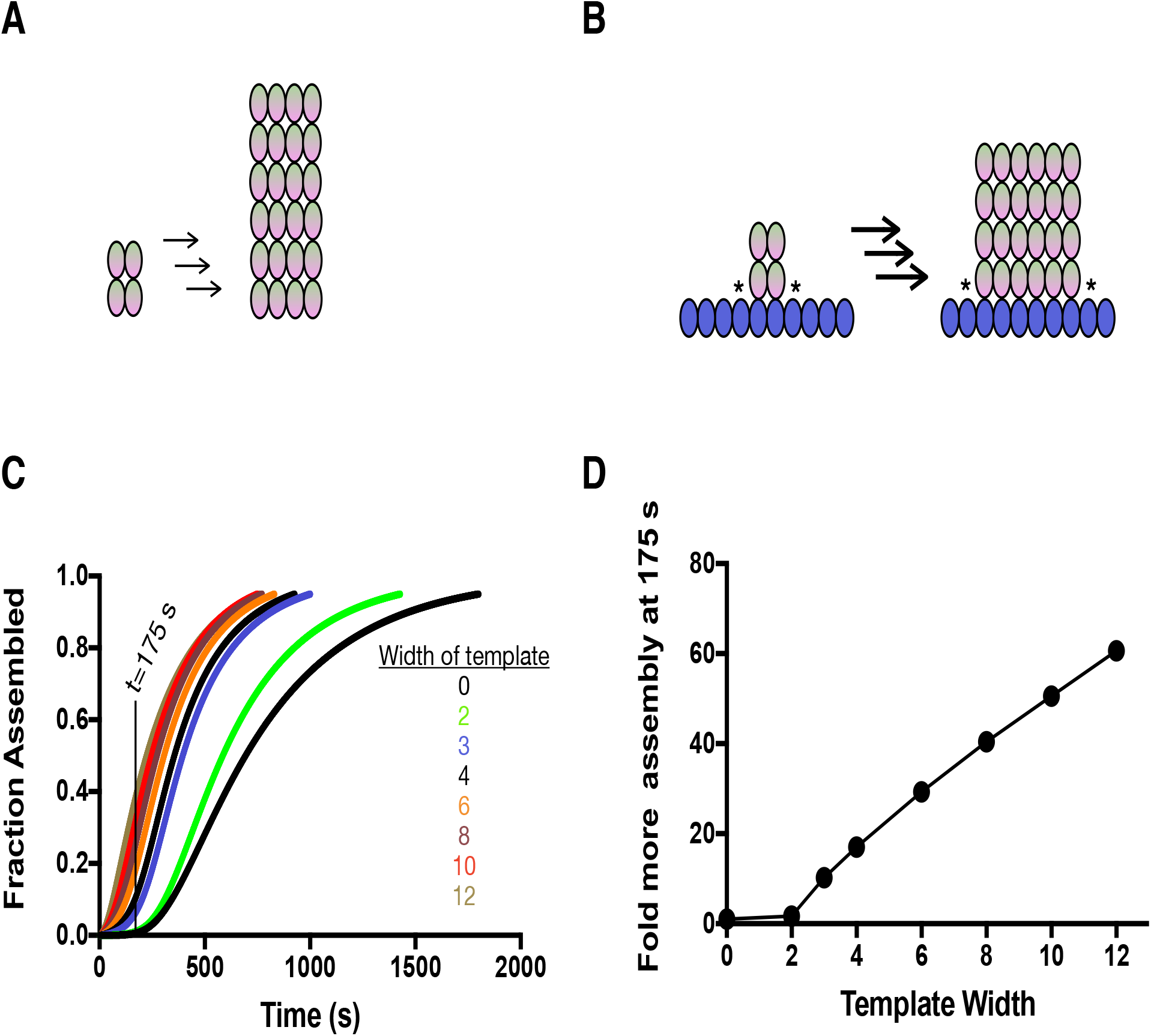
Implications for templating microtubule assembly **A** Cartoon illustrating an intermediate stage of spontaneous nucleation. 2×2 oligomers of αβ-tubulin (left) assemble as described into rectangular intermediates that tend to be taller than they are wide (middle) because of the barriers to lateral growth. Arrows indicate multiple layer additions, each one associated with a delay. **B** When αβ-tubulin adds to a γ-TuRC-like template, cartooned here as a row of blue ovals, the barrier to lateral growth is eliminated because ‘corner’ sites form at the border of the αβ-tubulin oligomer. This accelerates the accretion process (bolder arrows indicating faster transitions) and the resulting oligomers now tend to be wider than they are tall **C** Simulated nucleation curves in the presence of γ-tubulin templates of different width assuming γ:αβ interactions are weaker than αβ:αβ interactions. The total concentration of templating monomers is held constant. Templates that are only two subunits wide (bright green) show a modest effect, but wider templates are increasingly potent. The vertical line indicates the 175 s timepoint that we used for analysis in the next panel. **D** Quantification of the data from C. Wider templates are more potent than narrower ones. Templating the earliest step in nucleation (from 2-wide to 3-wide) yields the most dramatic decrease in T0.1. **E** Fraction of αβ-tubulin incorporated into microtubules at an early time (here, 175 s, see vertical line in panel C) plotted as a function of template width and using data from C. Relative assembly at a given time represents the fraction assembled for a given template divided by the fraction assembled in the untemplated control. The changes in nucleation kinetics can result in large changes in the overall degree of microtubule assembly at early times.

The idea that multiple reactions/transitions contribute to limit the overall rate of microtubule formation also suggests that strategies besides templating could enhance the rate of spontaneous assembly. The rates for layer additions depend on how fast tubulins ‘land’ on the growing sheet, so enhancing this rate should promote layer initiation and ‘zippering’. For example, XMAP215 family polymerases use an ‘accelerated delivery’ mechanism (Ayaz et al., 2014; Geyer et al., 2018) to make microtubules grow faster, and this same activity likely underlies the role for XMAP215 family proteins in microtubule nucleation (Gunzelmann et al., 2018; Popov et al., 2002; Roostalu et al., 2015; Thawani et al., 2018; Wieczorek et al., 2015). The efficiency of initiating new layers is primarily limited by the relatively weak longitudinal or lateral affinities, especially for the singleton tubulin that initiates each longitudinal layer. Thus, a factor that stabilizes these associations should also increase the rate at which layer transitions occur. Proteins like TPX2 and doublecortin bridge between neighboring subunits in the lattice (Fourniol et al., 2010; Zhang et al., 2017); this ‘molecular stapling’ may help explain their documented roles in microtubule “nucleation” (Moores et al., 2004; Roostalu et al., 2015). Finally, altering the allosteric properties of αβ-tubulin itself should also profoundly affect microtubule initiation. For example, and in keeping with predictions of our model, studies of an αβ-tubulin ‘conformation cycle’ mutation (Geyer et al., 2015) that enhances the strength of lateral interactions (Driver et al., 2017) (perhaps via reducing the cost of αβ-tubulin straightening), greatly accelerates microtubule formation. Similarly, small molecule compounds like taxol that stimulate microtubule initiation and growth may do so by promoting straightening.

### Conclusions and perspective

We described a relatively simple accretion model that explains spontaneous microtubule assembly kinetics in terms of subunit biochemistry and an ensemble of ‘rectangular’ intermediates. Whereas in typical ‘nucleation-elongation’ paradigms the scarcity of the transition-state-like nucleus is what limits the rate of polymer formation, our model is more akin to a process like crystal growth. Indeed, in our accretion model most of the steps toward polymer formation are energetically favorable but kinetically frustrated by the difficulty of initiating new layers; this difficulty decreases as layers become larger because more landing opportunities are presented. The accretion mechanism we describe is a direct consequence of the two-dimensional nature of the hollow, cylindrical microtubule lattice.

We did not consider the kinetic consequences of GTP hydrolysis within the nascent polymer in our model, because it was not possible to account for the stochastic hydrolysis within the context of our deterministic formalism. In keeping with that, our assay conditions were chosen to suppress microtubule catastrophe. We attempted to measure assembly curves in the presence of GMPCPP, a hydrolysis-resistant GTP analog, but obtained variable results because it was difficult to reliably eliminate contaminating amounts of microtubule seeds from the initial reaction mixtures. Ignoring GTPase activity represents a limitation of our model that we plan to address in future work. In general, GTPase activity should antagonize assembly by preferentially destabilizing intermediate oligomers, so the affinities we report may underestimate the ‘true’ affinities. We still expect that growth through accretion will represent the dominant pathway for assembly.

Finally, our model provides a unifying quantitative framework for understanding the complex effects of regulatory molecules. Indeed, the ‘zippering’ transitions between rectangular oligomers that dominate the initiation process also share many features in common with microtubule elongation. Thus, our model also explains why factors that regulate microtubule elongation and shrinking also regulate spontaneous microtubule assembly.

## Acknowledgements

This study was supported by grants from the National Science Foundation to LMR (MCB-1615938 and MCB-1054947) and from the National Institutes of Health to DAA (R01 GM031627, R35 GM118099 and P01 GM105537). DAA was supported in part by the Howard Hughes Medical Institute. We thank T. Davis and her colleagues for very helpful feedback on the manuscript.

## Author contributions

MM collected the microtubule assembly data and helped write the manuscript.

LMR constructed models, fit the data, and drafted the manuscript. DAA analyzed data, supervised the research, and helped write the manuscript.

## Methods

### Measurements of spontaneous assembly

The spontaneous assembly of phosphocellulose-purified porcine-brain tubulin was followed by turbidity at 350 nm basically as described previously (Gaskin et al., 1974; Voter and Erickson, 1984). Care was taken to remove microtubule seeds and inactive protein by cycling it through an additional polymerization/depolymerization step just before use: immediately prior to performing assays, tubulin at –80°C was rapidly thawed at 37°, placed on an ice-water slurry (0°C), and the buffer was adjusted to 80 mM K-PIPES, pH 6.8, 1mM EGTA, 4 mM MgCl_2_ and 1 mM GTP. After 5 min incubation, 0.5 volume of 37°C glycerol was mixed in and the tubulin was allowed to polymerize at 37°C for 20 min. Microtubules were pelleted through a 37°C, 60% glycerol cushion in 50 mM K-MES, pH 6.6, 5 mM MgSO_4_, 1 mM EGTA, 1 mM GTP in a TLA110 rotor (Beckman, Palo Alto, CA) at 80,000 rpm, 20 min. The pellets were resuspended at 37°C in assembly buffer (AB: 50 mM K-MES, pH6.6, 3.4 M glycerol, 5 mM MgSO_4_, 1 mM EGTA, 1 mM GTP) using a warm Potter homogenizer. The resuspended microtubules were then depolymerized on an ice-water slurry for 20 min and centrifuged in a TLA100.3 rotor at 100,000 rpm for 35 min at 2°C to remove any remaining polymerized tubulin. The supernatant from this cold spin was removed and used for the assembly reactions. The tubulin was kept at 0°C as reaction mixtures were prepared. A series of samples was prepared at different tubulin concentrations in AB and kept at 0°C in thin-walled, 0.5 ml plastic tubes (PCR tubes, Stratagene, Cedar Creek, TX). Samples were rapidly heated to 37°C directly in cuvettes pre-warmed to 37°C in a Pelltier cell changer (took 30 sec), in a custom built spectrophotometer. The A_350 nm_ was recorded approximately every 4 sec on 4 samples at a time until plateaus were reached. Recording on the spectrometer was started prior to transfer of tubulin to the cuvettes so that early lag-phase data were not missed. The time was adjusted to ignore the 30 sec heating step. Scaling analysis was performed using a customized program.

### Explicit simulations in Berkeley Madonna

We constructed a model in Berkeley Madonna (Marcoline et al., 2020) to simulate αβ-tubulin self-assembly. The first step in this process was to define the set of αβ-tubulin oligomers and how they are related to each other by gain/loss of an αβ-tubulin though a longitudinal, lateral, or longitudinal+lateral (‘corner’) interaction. Starting from the two-dimensional nature of the microtubule lattice, we built a list of all possible configurations for an αβ-tubulin oligomer of a given size. Fig. 1 illustrates how there are two kinds of possible dimers of αβ-tubulin (longitudinal and lateral), three kinds of trimers of αβ-tubulin, etc. Larger oligomers can have multiple configurations that are energetically equivalent. To minimize the number of species and reactions considered in the model, we used a single exemplar species to represent each set of n-mers with the same number of longitudinal and lateral contacts. For each exemplar species, population-weighted average rate constants were used to account for the different properties of each oligomer. Supplemental Figure 1 provides a worked example of this approximation for a tetramer of αβ-tubulin. We then used the ‘Chemical Reactions’ module in Berkeley Madonna (Marcoline et al., 2020) to build a set of rate equations corresponding to each possible transition. We did not pursue oligomers larger than 12-mers because the number of species and reactions became intractable. This model provides a way to analyze the rate and extent to which different species become populated for a given choice of the adjustable parameters.

### Derivation of layer transition rates and validation in Berkeley Madonna

The explicit, monomer-based simulations in Berkeley Madonna (Marcoline et al., 2020) indicated that a simpler reaction scheme that only considered transitions between rectangular oligomers. We used the steady state approximation to derive effective rate constants for these transitions, as described below.

We will illustrate the derivation of layer-based transition rate constants using a specific example. Consider the layer-based reaction associated with adding a longitudinal layer onto an oligomer that is, say, n tubulins tall and 4 tubulins wide (in other words, the net reaction is (n,4) <--> (n+1,4). To simplify the notation, we will call the starting species **4** and the final species **4’**; the intermediates with one, two, and three tubulins on the layer will be called 4a, 4b, and 4c respectively. The individual reactions are:

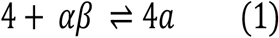

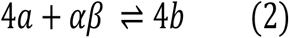

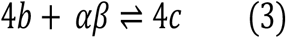

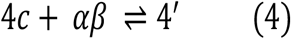

The associated rate equations for these reactions, which account for the production and loss of intermediate species and also the fact that different configurations present different numbers of binding sites (Sup. Fig. 8), are:

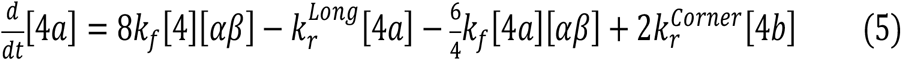

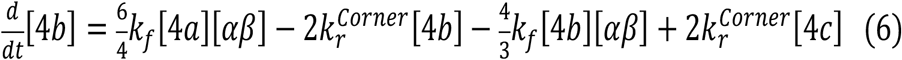

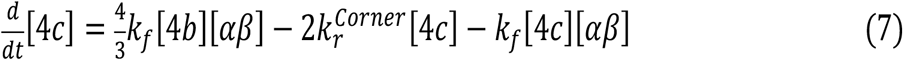

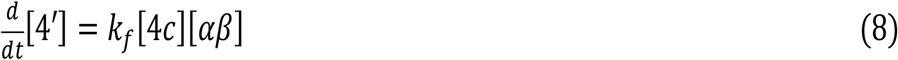

Starting from the last intermediate 4c, we apply the steady-state approximation to the intermediate species to obtain an expression for the net rate of producing 4’ from 4. Assuming that the concentration of 4c does not change with time, its time derivative is zero, so:

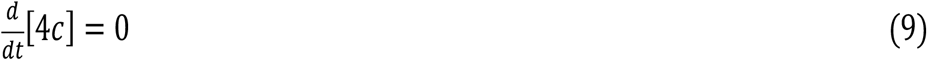

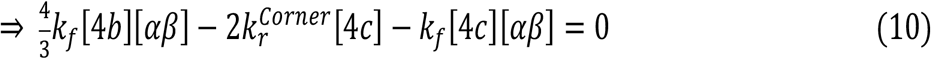

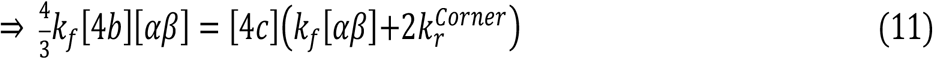

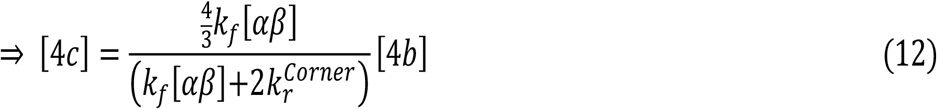

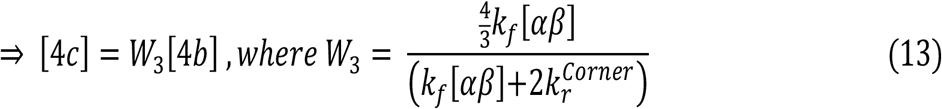

This yields an expression for the concentration of intermediate 4c in terms of its predecessor 4b. The multiplicative factor W3 is given by the pseudo-first-order rate constant for forming 4c divided by the sum of the pseudo-first-order rate constants for losing 4c (in the forward or reverse direction).

This process can be applied again to obtain an expression for the concentration of intermediate 4b in terms of its predecessor 4a:

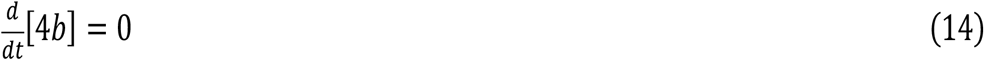

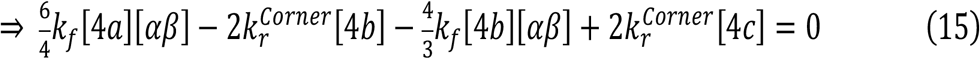

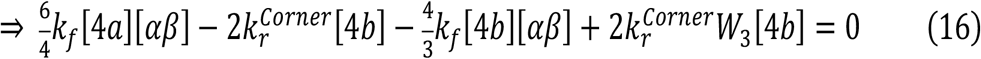

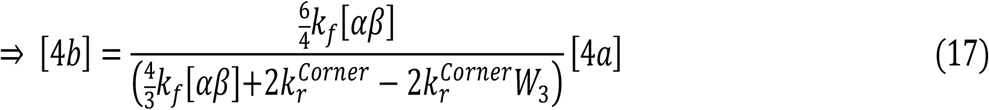

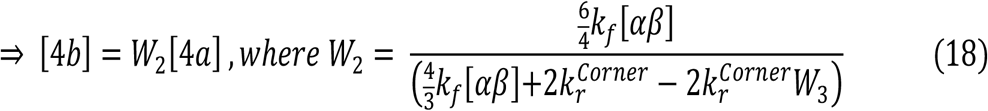

Where in (16) we eliminated [4c] by substituting W3[4b], and in (17) we simply grouped terms in [4a] and [4b]. The weighting term W2 is again given by a similar ratio as for W3, but with an additional term that includes W3.

Applying the same procedure, we can obtain an expression for the concentration of the first intermediate 4a in terms of its predecessor, the rectangular species 4:

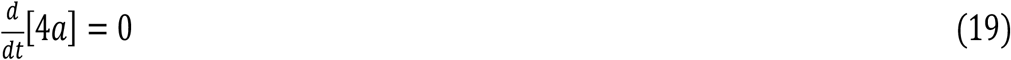

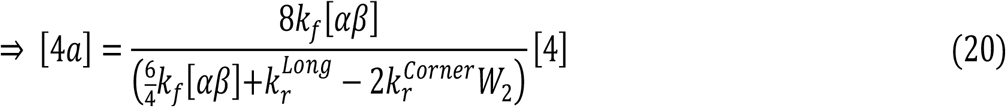

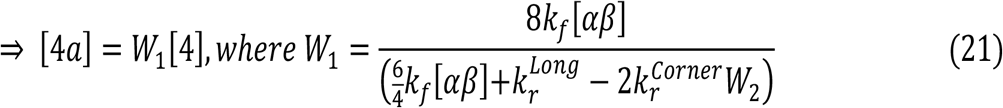

Finally, since the rate of finishing the layer is given by:

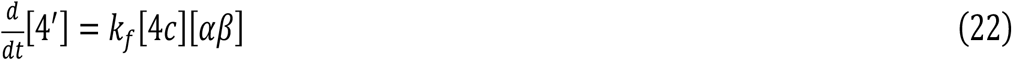

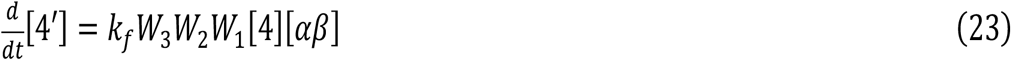

Where in (23) we used (13) to eliminate [4c] for a term involving [4b], (18) to eliminate [4b] for a term involving [4a], and (21) to eliminate [4a] for a term involving [4]. This yields an apparent rate constant for adding a layer, the terms of which only involve the fundamental rate constants and multiplicative factors that account for the number of ways a particular reaction can happen. In the case presented there are three such terms because the layer contains 4 tubulins. Because k_r_^Corner^ is small compared to k_f_[αβ], is can be neglected in the denominator and so all weights excepting the first one will be numbers between 1 and 2. W_1_ contains k_r_^Long^ in the denomimator, which in general is large compared to k_f_[αβ], so W_1_ tends to be a small number that reflects the difficulty of initiating a new layer. The general procedure is formulaic and can be applied to any size layer.

The effective equilibrium ratio for the layer addition is given by

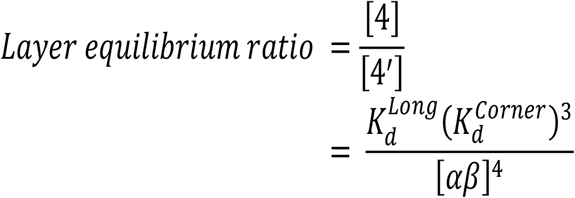

Finally, we compute an effective rate constant for layer loss by multiplying the layer equilibrium ratio and the forward layer rate constant:

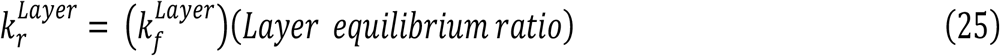

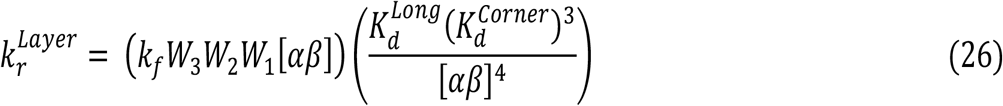

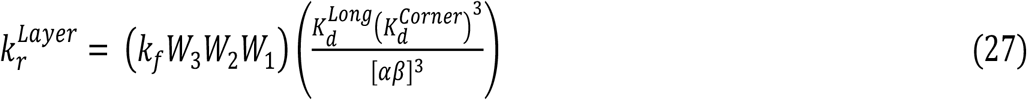

The preceding derivation was performed for the specific case of adding a longitudinal layer that is four subunits wide. For layers of size N, a similar logic applies with the following differences: (i) the kinetic weighting factors for the first through the (N-1)^th^ subunit additions change to reflect the different number of configurations (Sup. Fig. 8), and (ii) there is a different number of multiplicative factors (*W_i_s*).

To determine if this reduced representation could accurately recapitulate the timecourses for rectangular species in the explicit model, we constructed a new model in Berkeley Madonna (Marcoline et al., 2020) (also truncated at 12-mers) that considered monomer-based reactions to form the first rectangular intermediate, a 2×2 tetramer of αβ-tubulin, but that only considered layer transitions for larger species, using the layer rate laws described above. We input expressions for the layer transition rates as derived above, and ran the simulations using the same biochemical parameters used in the explicit simulations. The layer model faithfully reproduced the results from the explicit simulations (Fig. 1).

### Layer-based simulations of microtubule self-assembly

To enable simulations of arbitrarily large oligomers, we wrote a custom computer program that effectively performs the same calculations of Berkeley Madonna but in a more automated way that makes it easier to incorporate a much larger number of species and reactions. The program automates the generation of the list of possible rectangular oligomers [(2,2), (3,2), (2,3), etc.] and the recursive calculation of rate constants for transitions between them (as explained above and in Sup. Fig. 8). The program then integrates the set of transition rate laws using finite-difference methods. At each time point, the program uses longitudinal and lateral affinities along with the concentration of ‘free’ tubulin (here free means the population not incorporated into rectangular oligomers (2,2) or larger) to calculate the concentration of free tubulin ‘monomers’ and of longitudinal and lateral dimers, and trimers assuming these species are all in rapid equilibrium (see box in Fig. 1E); using these quantities, the program then takes a small step forward in time, updating quantities of all rectangular oligomers according to the rate laws and the layer rate constants. This results in an updated set of concentrations for the rectangular oligomers (and hence of free tubulin), and the process is repeated. To account for microtubule formation, we simply converted species that became 13-protofilaments wide into microtubules, and we assumed that microtubules elongated by monomer addition into a ‘corner’ site. We did not explicitly consider the seam or the variable configuration of the microtubule end, and for simplicity we assumed plus- and minus-ends elongated with identical kinetics. We used a downhill simplex algorithm to fit the simulated timecourses to the experimental ones, using fraction of tubulin in rectangular oligomers as a proxy for the normalized assembly data.

### Derivation of the equilibrium constant for lateral association of a pre-formed longitudinal dimer

To implement the dimer pathway (Fig. 2 and Sup. Fig. 4), we assume that the on-rate constant for adding a dimer is the same as for adding a monomer. Given the on-rate constant, we obtain the off-rate constant (for the initiating dimer to dissociate) using the equilibrium constant for the dimer binding. We show here that this new equilibrium constant can be calculated purely in terms of ‘monomer’ properties. First, we write the dissociation constants in terms of free energies of association, as follows:

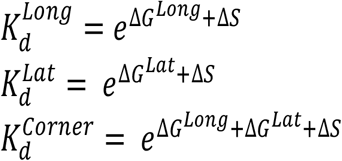

where we have separated out the entropic cost of subunit immobilization (loss of rotational and translational degrees of freedom) as ΔS as has been done previously (e.g. (Erickson and Pantaloni, 1981; VanBuren et al., 2002)). The reason for doing this separation is that lateral association of a pre-formed longitudinal dimer should have the same entropic ‘cost’ as for association of a monomer (because the cost of losing rotational and translational degrees of freedom is already built in to the longitudinal affinity). The K_d_ values are the fitted parameters in the model.

What we wish to calculate is the dissociation constant for two simultaneous lateral contacts:

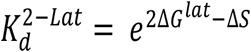

We prove here that:

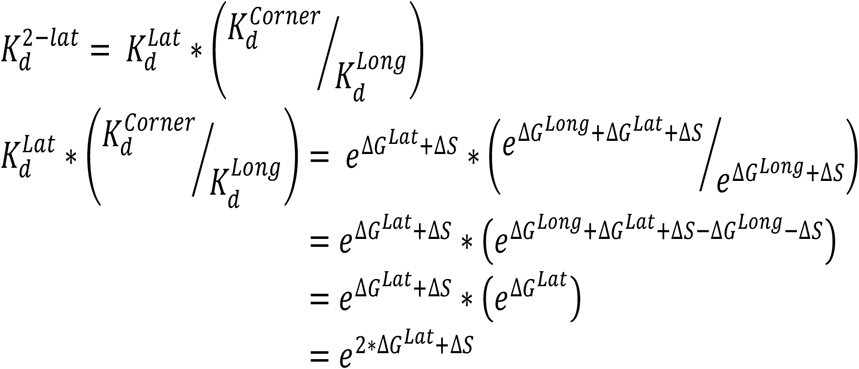

### Simulating the effects of template-like nucleators

To investigate the effects of template-like nucleators in the model, we modified the simulations to include an additional template species. We assumed a constant total concentration of ‘monomers’ that were distributed into templating oligomers of different widths. This assumption means that progressively wider oligomers are present in the simulations at progressively lower concentrations; for example, for a given concentration of templating monomers, there will be half as many 4-wide oligomers as 2-wide, and so on. For calculating the rate constants for the first layer addition onto the template, we assumed that only one longitudinal interface of this templating oligomer (the plus end) was active for nucleation.

In initial runs wherein interactions between γ- and αβ-tubulin were assumed to be of identical affinity, we observed potent nucleation even from templates that were only two subunits wide (Supplemental Fig. 7). Because two-wide nucleators like the γ-TuSC are much less potent than 12-14-wide oligomers like the γ-TuRC (Kollman et al., 2015; Oegema et al., 1999), we also performed simulations with 10-fold weaker interactions between γ- and αβ-tubulin than for αβ:αβ interactions (Fig. 6)

---

**Supplemental Figure 1.**
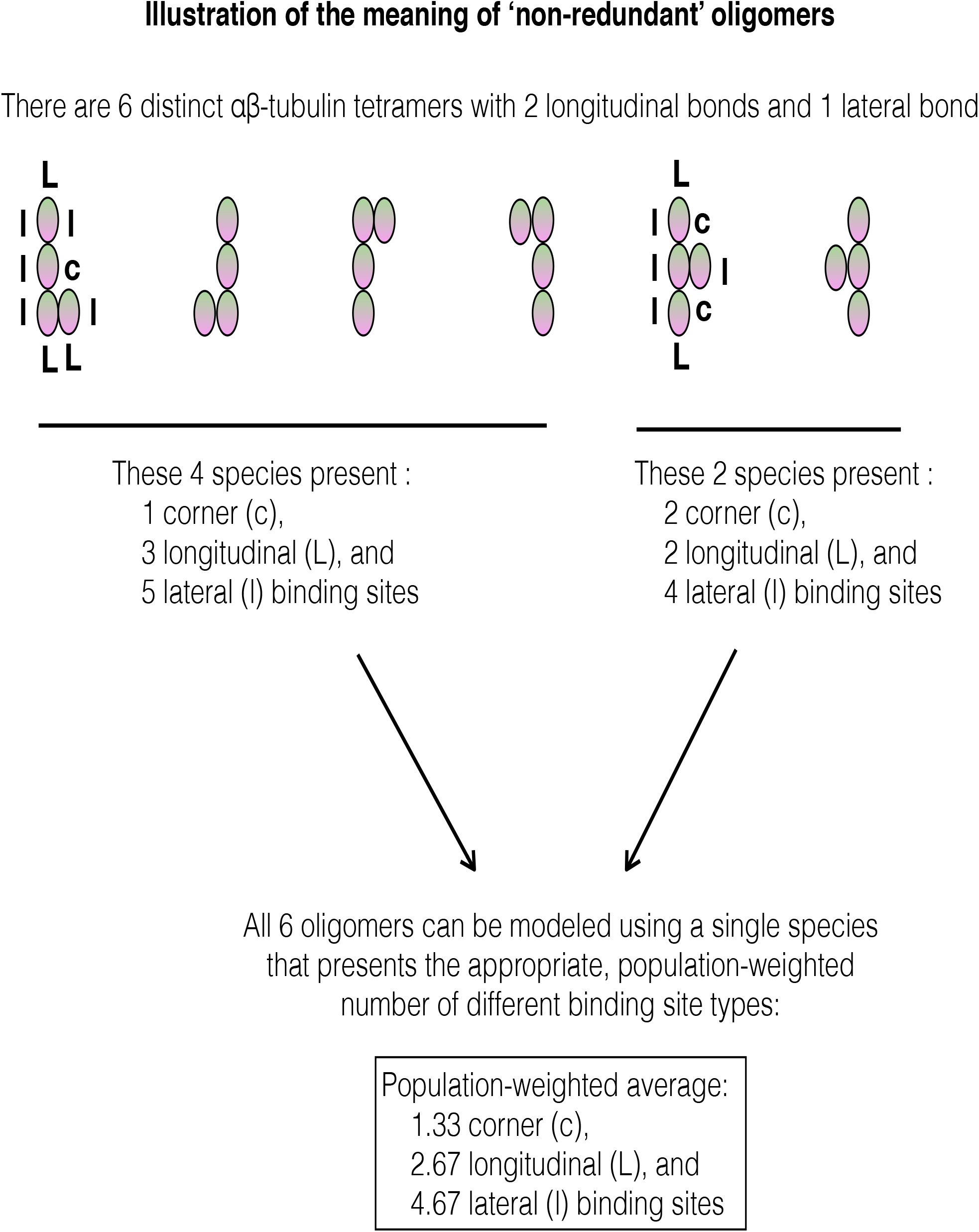
Illustration of the concept and implementation of ‘non-redundant’ oligomers. Representative configurations of an αβ-tubulin tetramer containing two longitudinal and one lateral bond are indicated. These are indistinguishable from the standpoint of binding energy, but differ in terms of the number and type of binding sites they present. Our algorithm simplifies these six species into one and uses population weighted rate constants to capture the ‘average’ number and type of binding sites presented.

**Supplemental Figure 2.**
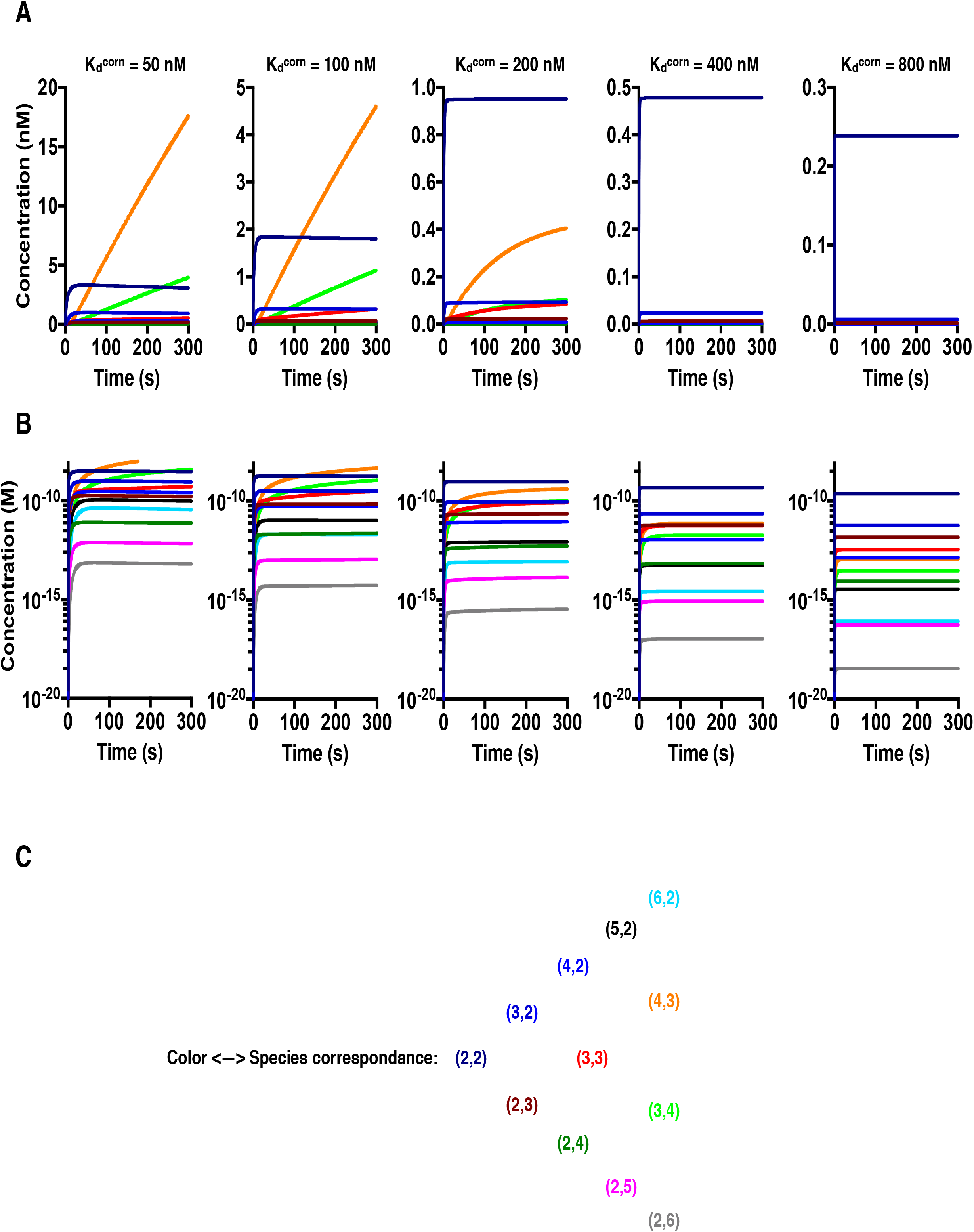
In the explicit model, relatively tight corner affinity is required for appreciable assembly into larger oligomers. **A** Plots of the concentration of ‘rectangular’ oligomers vs time, for different values of the corner affinity (color coding is given in panel C and is identical to that used in Fig. 2B). Note different y-axis ranges for different plots. At corner affinity of 400 nM and weaker, assembly into species larger than a tetramer of αβ-tubulin (dark blue curve) is negligeable. **B** As in A, but using a log-scale y-axis to show the dramatic fall-off in the concentration of larger species. **C** Legend showing the correspondence between color and oligomer dimensions.

**Supplemental Figure 3.**
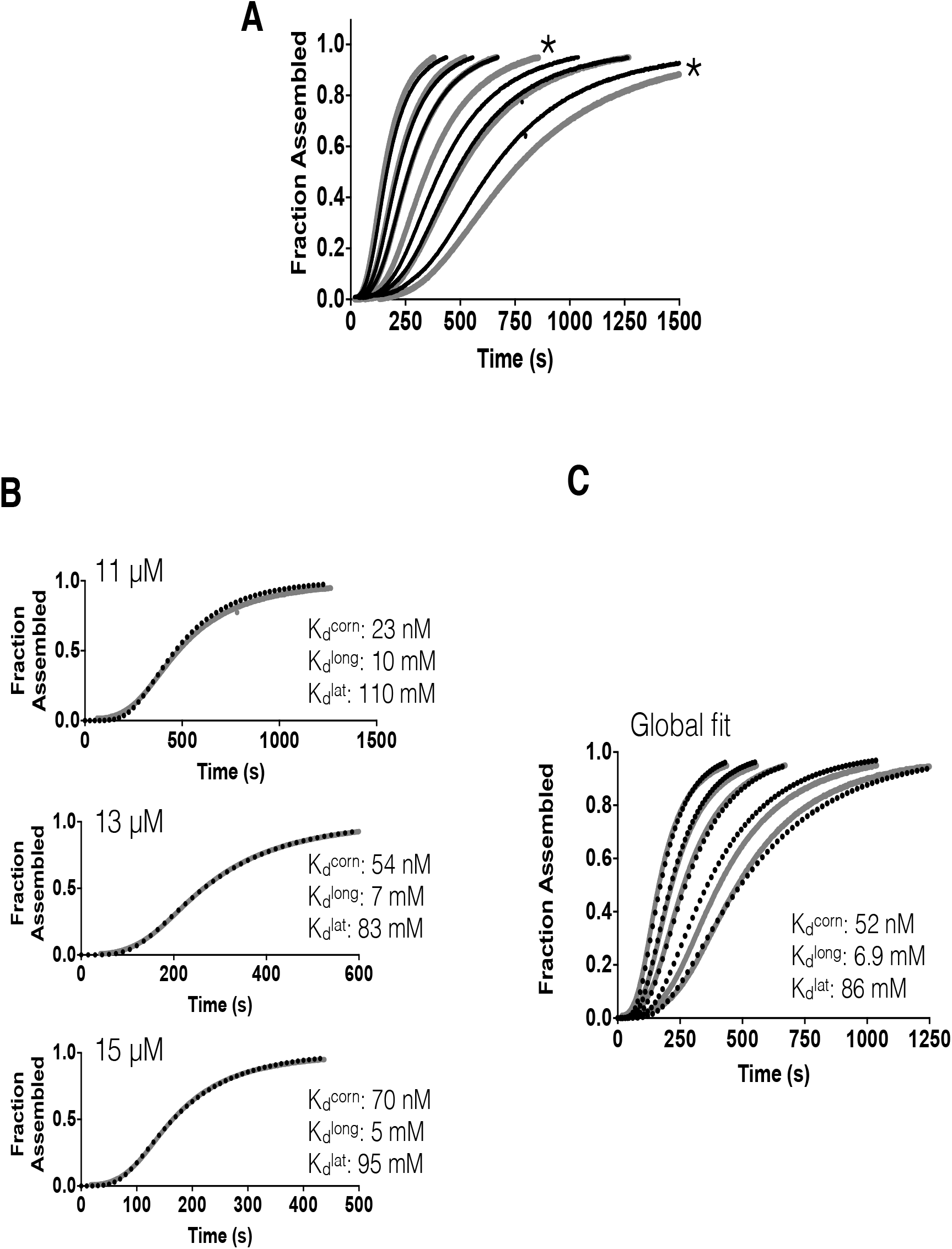
Fit to an independent dataset **A** Two independent data sets. The set plotted in grey is the one analyzed in the main text. The set in black was collected independently. Tubulin concentrations were 10, 11, 12, 13, 14, 15 μM for each. Most of the curves agree very closely, with the exception of two outlier curves (10 and 12 μM, indicated with # and *, respectively). The 10 μM curve was omitted from panel C. **B** Fits of the layer model (black dotted line) to individual assembly curves (solid grey line) at different concentrations. Fits to individual curves are good. Inset text shows the resulting longitudinal, lateral, and ‘corner’ (simultaneous longitudinal and lateral interactions) affinities, which are within 10% of those obtained in the main text (see Fig. 2B). **C** Global fit of the model (black dashed lines) to the alternate set of assembly data (grey lines). The model underestimates the concentration-dependence of the progress curves.

**Supplemental Figure 4.**
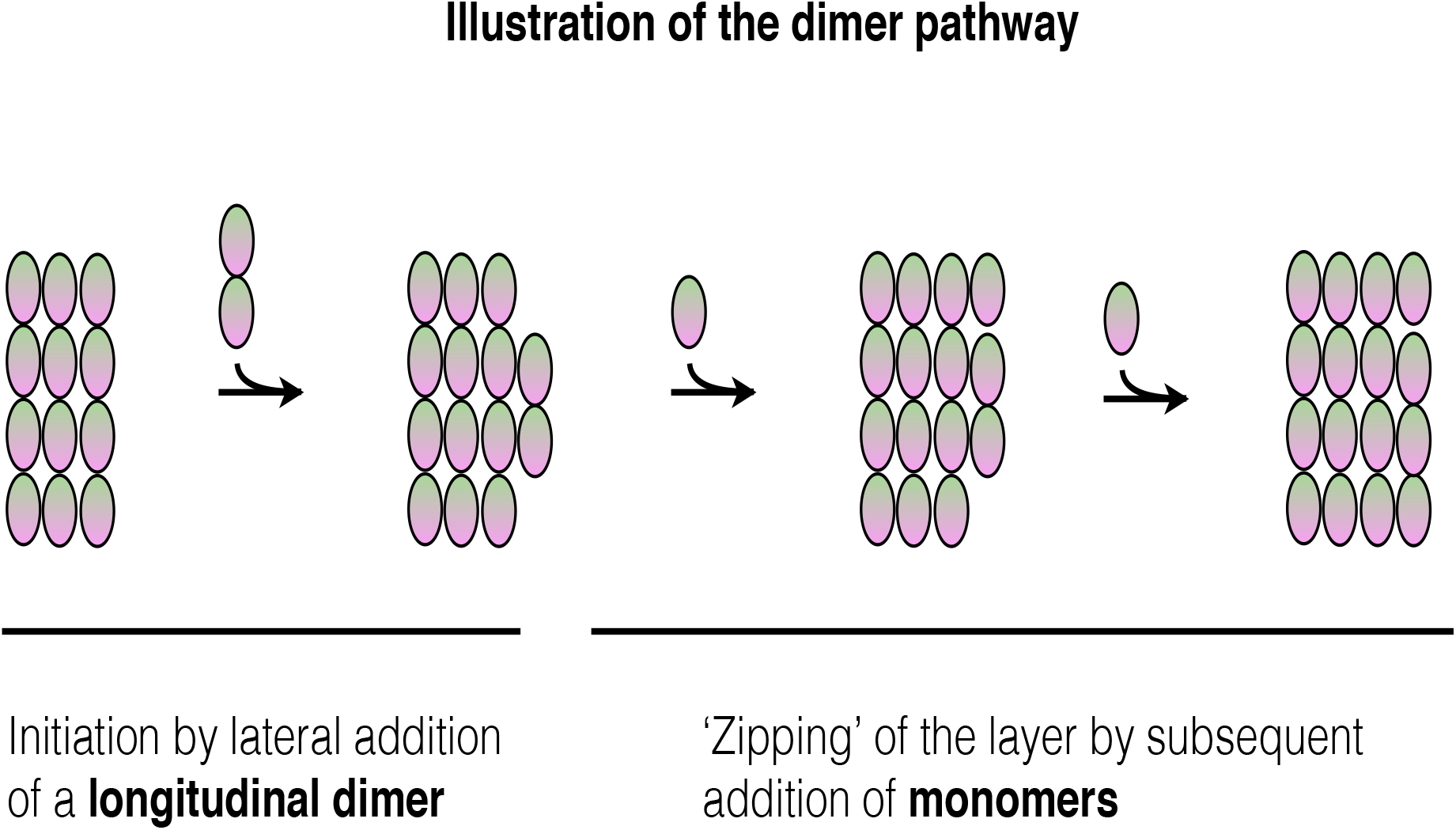
Illustration of the dimer pathway. Cartoon sequence of steps for adding a new lateral layer. In the dimer pathway, a new lateral layer can be initiated by association of a pre-formed longitudinal dimer (left). Because the concentration of these longitudinal dimers of αβ-tubulin is much less than that of αβ-tubulin ‘monomers’, subsequent ‘zipping’ of the layer occurs by monomer addition. See Methods for a derivation of the equilibrium binding constant for the initiating assocition.

**Supplemental Figure 5.**
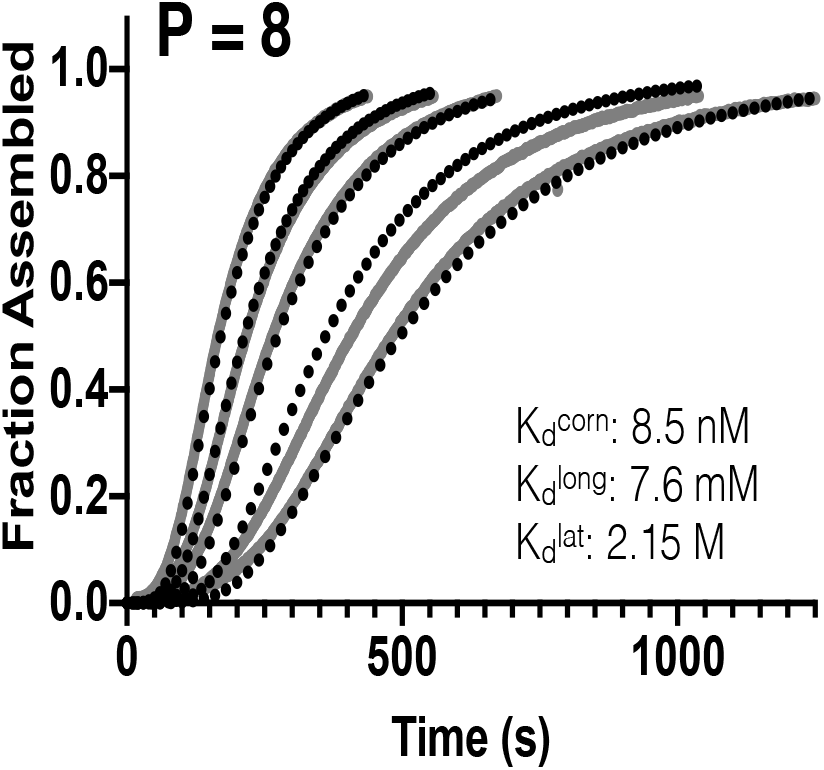
Fit of the dimer+penalty model to an independent dataset Global fit of the dimer+penalty model (black dotted lines) to the alternate set of assembly data (grey lines). The global fit is good (with the exception of the 12 μM outlier curve noted in Sup. Fig. 3). The fitted parameters are close to those obtained from fitting to the original dataset, but for this dataset the corner affinity is ~1.4-fold stronger and the longitudinal affinity is ~1.3-fold weaker.

**Supplemental Figure 6.**
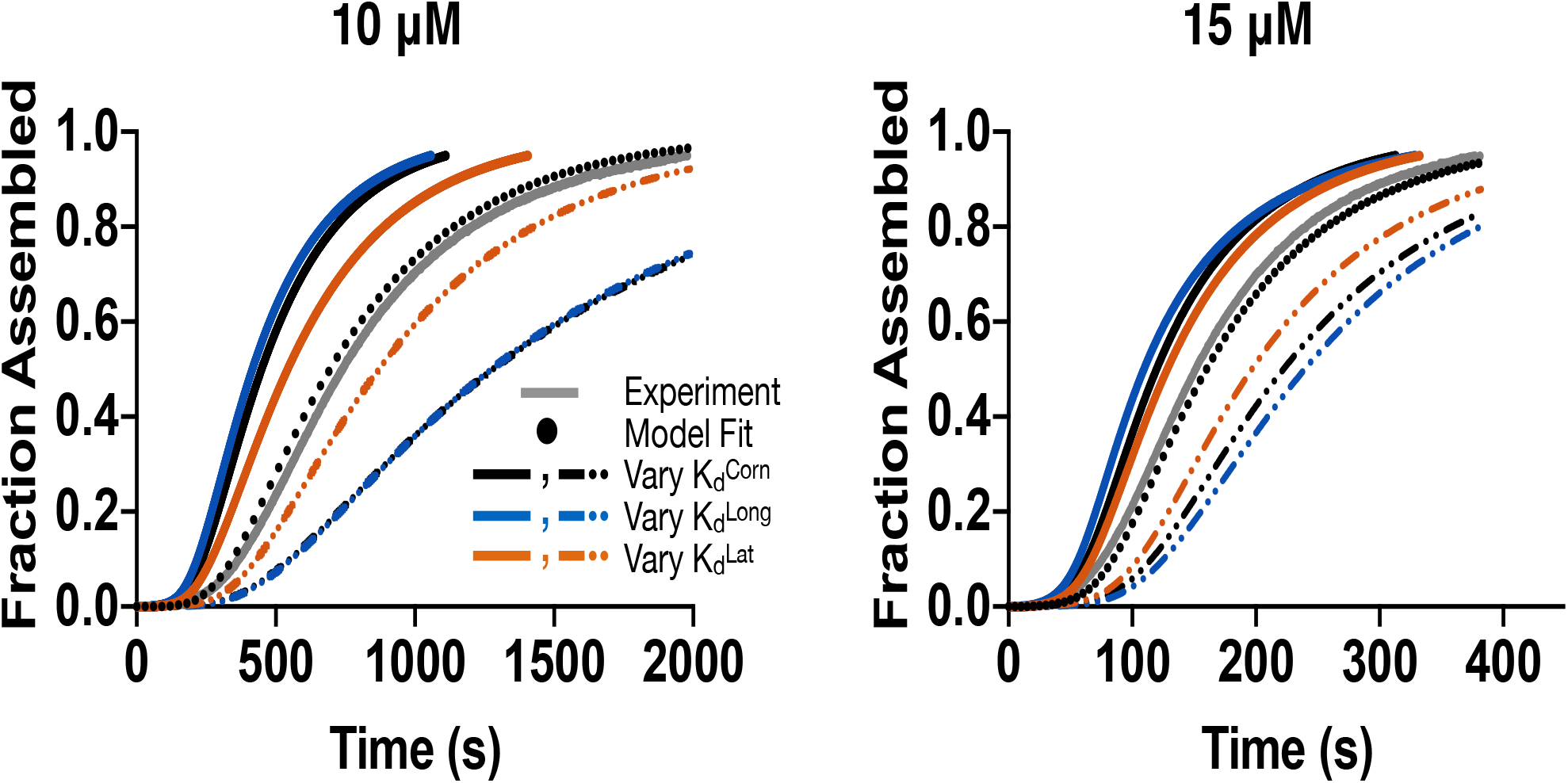
Sensitivity of model predictions to variations in the underlying parameters. The dimer+penalty model was run with individual affinities made stronger (solid lines) or weaker (dashed lines) by a factor of 1.5. Two different concentrations are illustrated. The model predictions are comparably sensitive to variation in corner (black) and longitudinal (blue) affinities, and less sensitive to variation in the lateral affinity (orange).

**Supplemental Figure 7.**
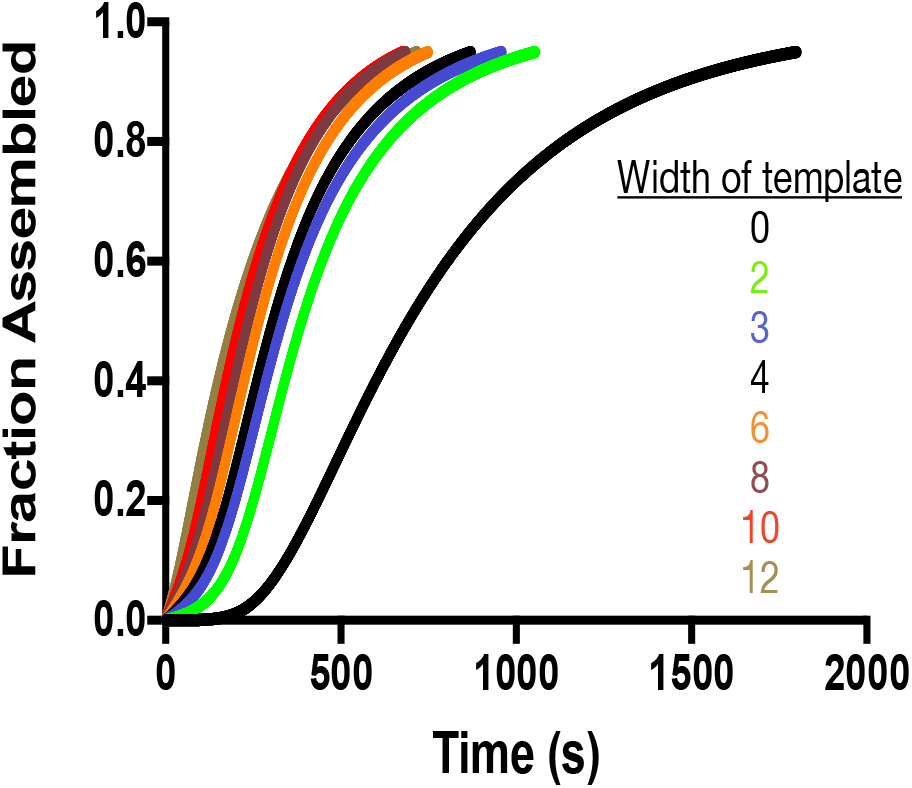
Simulations of templated microtubule assembly. Simulated nucleation curves in the presence of γ-tubulin templates of different width assuming γ:αβ interactions are the same strength as αβ:αβ interactions. Compare to Fig. 6. The total concentration of templating monomers is held constant. Templates that are only two subunits wide (bright green) show much stronger nucleation than when we assumed weaker γ:αβ interactions (compare green curve to the one in Fig. 6).

**Supplemental Figure 8.**
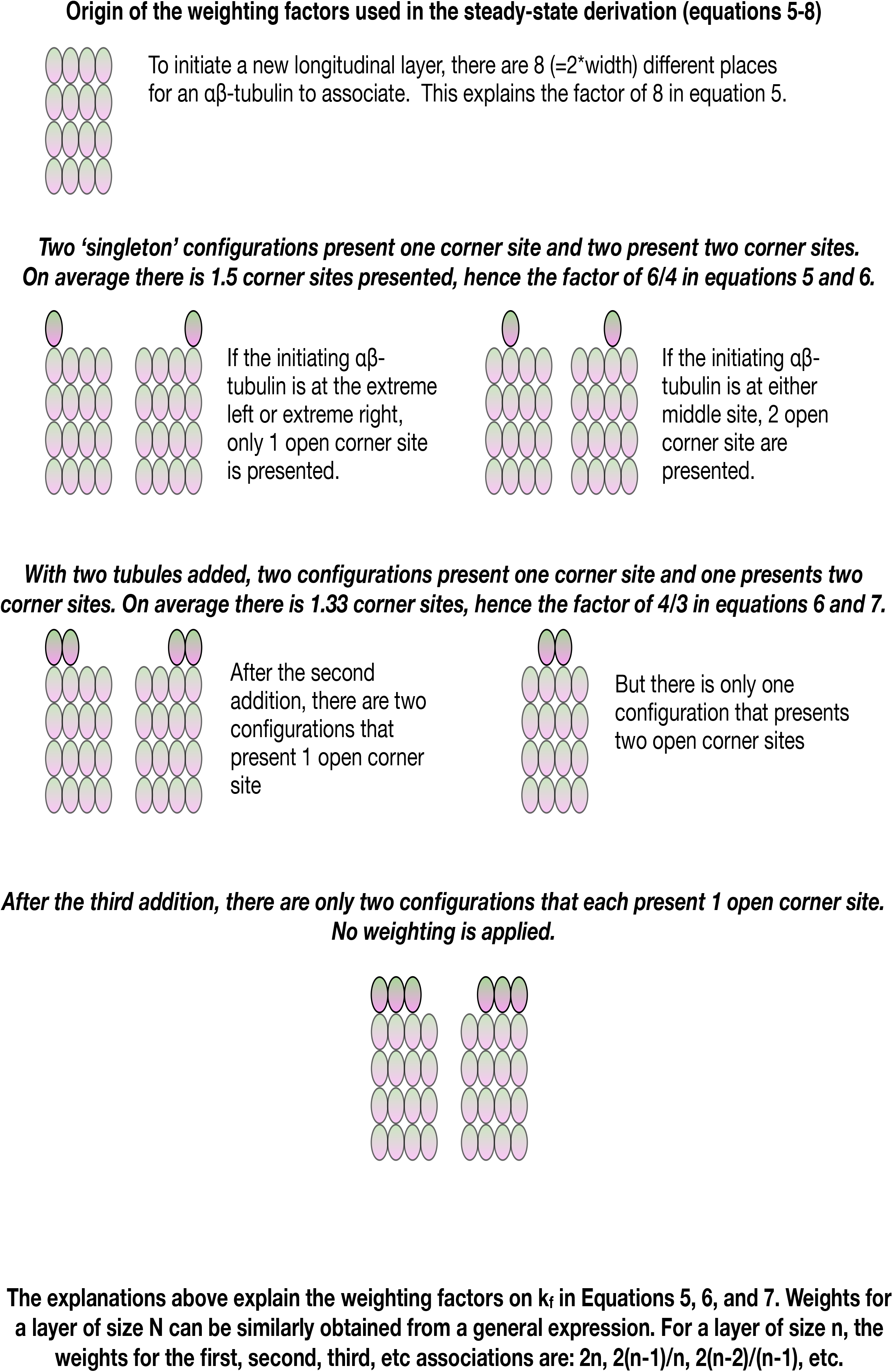
Derivation of kinetic weighting factors Cartoons illustrating the sequence of intermediates for a ‘layer addition’, annotated to indicate the origin of the weighting factors applied in equations 5–8. Illustrations are made for the specific case of a layer of size 4, but the general expression for the coefficients is provided at the bottom of the figure.

